# Subiculum – BNST Structural Connectivity in Humans and Macaques

**DOI:** 10.1101/2021.11.11.468209

**Authors:** Samuel C. Berry, Andrew D. Lawrence, Thomas M. Lancaster, Chiara M. Casella, John P. Aggleton, Mark Postans

## Abstract

Invasive tract-tracing studies in rodents implicate a direct connection between the subiculum and bed nucleus of the stria terminalis (BNST) as a key component of neural pathways mediating hippocampal regulation of the Hypothalamic-Pituitary-Adrenal (HPA) axis. A clear characterisation of the connections linking the subiculum and BNST in humans and non-human primates is lacking. To address this, we first delineated the projections from the subiculum to the BNST using anterograde tracers injected into macaque monkeys, revealing evidence for a monosynaptic subiculum-BNST projection involving the fornix. Second, we used in vivo diffusion MRI tractography in macaques and humans to demonstrate substantial subiculum complex connectivity to the BNST in both species. This connection was primarily mediated through the fornix, with additional connectivity via the amygdala, consistent with rodent anatomy. Third, utilising the twin-based nature of our human sample, we found that microstructural properties of these tracts are moderately heritable (h^2^ ∼ 0.5). In a final analysis, we found no evidence of any significant association between subiculum complex-BNST tract microstructure and indices of perceived stress/dispositional negativity and alcohol use, derived from principal component analysis decomposition of self-report data. We did, however, find subiculum complex-BNST tract microstructure associations with BMI, age, and sex. Our findings address a key translational gap in our knowledge of the neurocircuitry regulating stress.

## 1. Introduction

Though widely studied for its role in episodic memory and spatial navigation, the hippocampal formation also plays an important role in emotion, motivation, stress, fear and anxiety (for a review see Murray et al., 2017). In particular, structural connections between the main output region of the hippocampus, the subiculum (Aggleton & Christiansen, 2015; Böhm et al., 2018; Kishi et al., 2000), and the hypothalamic paraventricular nucleus (PVN) are implicated in control of the stress response via modulation of the hypothalamic-pituitary - adrenal (HPA) axis (Herman et al., 2003, 2020). Altered hippocampal regulation of the stress axis has been linked to various forms of psychopathology (Belleau et al., 2019; Herman et al., 2020). Studies in the rat suggest that subiculum modulation of the HPA axis is routed principally via structural connections to a key intermediatory structure, the bed nucleus of the stria terminalis (BNST) (for reviews see Herman et al., 2020; Radley & Johnson, 2018).

The BNST is a small grey matter structure (∼190mm^3^ in humans) situated within the basal forebrain (Alheid & Heimer, 1988; Johnston, 1923; Lebow & Chen, 2016). Commonly grouped with the central nucleus of the amygdala, due to dense connectivity via the stria terminalis fibre bundle and similarities in cytoarchitecture and function, the BNST forms a key part of the ‘Extended Amygdala’ (Alheid et al., 1998; Alheid & Heimer, 1988; Johnston, 1923). The BNST itself has been associated with a variety of processes, including anxiety, fear, substance abuse, reward, sexual motivation, and appetite (Avery et al., 2016; Fox & Shackman, 2019; Lebow & Chen, 2016). Although numerous rat studies implicate the BNST as a crucial intermediary structure en route from the subiculum to the PVN, descriptions of subiculum-BNST structural connectivity in the monkey and human literature are conspicuous by their absence (Aggleton & Christiansen, 2015; Böhm et al., 2018; Fox & Shackman, 2019; Lebow & Chen, 2016; Roberts & Clarke, 2019). Therefore, the present study aims to bridge this translational gap by characterising, via ex-vivo anatomic tract-tracing and in-vivo diffusion MRI tractography, this overlooked yet potentially key stress-regulatory pathway in both humans and macaques.

### 1.1 Subiculum - BNST Connectivity in Rodents

Invasive tract-tracing has demonstrated that the subiculum-BNST connection in rats and mice emanates from the ventral subiculum and travels via the fornix, or through a route involving the amygdala, to the BNST (Canteras & Swanson, 1992; Cullinan et al., 1993; Dong et al., 2001; Herman et al., 2020; Howell et al., 1991; Kishi et al., 2000; McDonald et al., 1999; Radley & Sawchenko, 2011; Shin et al., 2008; Swanson & Cowan, 1977).

In a key series of experiments, Cullinan, Herman, & Watson (1993) used the anterograde neuronal tracer Phaseolus vulgaris-leucoagglutinin (PHA-L) to reveal dense fornical and amygdala/ stria terminalis projections of ventral subiculum axons to the BNST in rats (Cullinan et al., 1993). They further used a mix of anterograde tracer PHA-L and retrograde tracer Flouro-gold to show that ventral subiculum neurons project to cells within the BNST that in turn project to the PVN (Cullinan et al., 1993). The researchers demonstrated that this takes place via a polysynaptic glucocorticoid - GABAergic feedback inhibition system, with the subiculum stimulating the BNST, which then sends inhibitory signals to the PVN (Cullinan et al., 1993). Whilst it was previously known that the subiculum is involved in top-down regulation of the HPA-axis (e.g., Jacobson & Sapolsky, 1991), this study demonstrated that the ventral subiculum largely relies on the BNST as an intermediary structure to do so. These findings, and others since, have seen the BNST come to be understood as an important relay, or cortical processing hub (Radley & Johnson, 2018), integrating information from the hippocampus and other regions (Radley & Sawchenko, 2011) before regulating HPA axis activity (Cullinan et al., 1993; Herman et al., 2020; Lebow & Chen, 2016). Research in rats suggest a specific role of the ventral subiculum – BNST pathway in response to acute stressors involving exteroceptive triggers (e.g., stimuli connected with novel environments), but not interoceptive cues (e.g., hypoxia) (Herman et al., 1998, 2020; Mueller et al., 2004; Radley & Sawchenko, 2011). However, the precise intricacies of how this pathway is related to the stress response is likely to be complicated by the heterogenous and complex structure of the BNST, with researchers demonstrating sometimes opposing functions of different BNST sub-regions on the PVN (Choi, Evanson, et al., 2008; Choi, Furay, et al., 2008; Herman et al., 2020; Myers et al., 2012).

### 1.2 Subiculum – BNST Connectivity in Monkeys

Subiculum-BNST connectivity in monkeys has been far less extensively studied. Probably the first indications of a corresponding connection in monkeys were published in studies that used a mix of electronic stimulation and lesion-degeneration methods to investigate hippocampal outputs in squirrel monkeys (Poletti et al., 1973; Poletti & Creswell, 1977; Poletti & Sujatanond, 1980). This research uncovered evidence for a light subiculum connection to the BNST (Poletti et al., 1973; Poletti & Creswell, 1977; Poletti & Sujatanond, 1980). In agreement with subsequent work in the rat (e.g., Cullinan et al., 1993) Poletti and colleagues also reported that hippocampal-BNST connectivity was not completely reliant on the fornix, but rather that the BNST receives additional hippocampal influences; likely involving the stria terminalis and/or the amygdalofugal pathway (Morrison & Poletti, 1980; Poletti et al., 1973; Poletti & Creswell, 1977; Poletti & Sujatanond, 1980). More recently, a ventral subiculum connection to the BNST has also been reported in the primate-like tree shrew (Ni et al., 2016).

### 1.3 Subiculum – BNST Connectivity in Humans

Diffusion MRI (dMRI) permits a non-invasive macro-scale estimate of white matter tracts in the brain by using the local restriction of water diffusion to infer the presence and orientation of white matter tracts (Jbabdi & Behrens, 2013). Researchers have demonstrated that pathways reconstructed in the human brain using dMRI often closely resemble those seen using monkey tract-tracing methods (Mars et al., 2018; Schmahmann et al., 2007), with an estimated 90% of ‘ground truth’ tracts being detectable (Maier-Hein et al., 2017). Owing to its small size and the relatively recent introduction of protocols for delineating the structure in the human brain, studies examining structural connections that include the BNST are rare. An exception (Avery et al., 2014) reported evidence of BNST structural connectivity to a variety of basal ganglia and limbic regions, including the hippocampus (Avery et al., 2014). Although only investigating the hippocampus as a single structure, an examination of the heat-map of connectivity strength within the hippocampus suggests that the subiculum is the principal driver of this connection (Avery et al., 2014, Figure 4).

As well as reconstructing tracts, dMRI estimates of water diffusion can be used to infer properties of white matter microstructure. For instance, fractional anisotropy (FA) can be used to estimate axonal myelination, axonal diameter, or fibre density (Basser, 1997; Dennis et al., 2021). Several human neuroimaging studies have reported associations, both phenotypic and genetic, between dMRI microstructure measures and human traits (e.g., Bauer et al., 2016; Cox et al., 2016; Gray et al., 2020; Lu et al., 2018; Rutten-Jacobs et al., 2018). Given rodent evidence implicating the subiculum– BNST connection in HPA-axis modulation (Herman et al., 2020), white matter microstructural properties of this tract may be related to traits and behaviours associated with stress reactivity.

Stressor reactivity is hypothesised to play a key role in dispositional negativity (Hur et al., 2019; Shackman et al., 2016). This term refers to individual differences in the propensity to experience and express more frequent, intense, or enduring negative affect (Hur et al., 2019; Shackman et al., 2016). Dispositional negativity is relatively stable across the lifespan and is predictive of numerous negative life outcomes, including elevated risk for the development of stress-sensitive psychiatric diseases; such as anxiety or depressive disorders (for reviews see; Hur et al., 2019; Shackman et al., 2016). Importantly, researchers have also connected the stress-response with potentially addictive behaviours, including alcohol use (Herman, 2012; Herman et al., 2020; Volkow et al., 2016). Evidence from both preclinical and human subjects have linked the hippocampus and the BNST to dispositional negativity and addictive behaviours (Hur et al., 2018; Lebow & Chen, 2016; Mira et al., 2020; Shackman et al., 2016). Alcohol consumption in particular has been shown to alter stress processing via drug-induced changes to PVN projecting limbic regions, including alterations to BNST and subiculum receptor signalling (Bach et al., 2021; Centanni et al., 2019; Haun et al., 2020; Mira et al., 2020). However, relationships between the microstructural properties of this putative subiculum-BNST pathway and dispositional negativity / alcohol use in humans are currently unknown.

### 1.4 Study Aims

This research will examine: 1) The evidence for a structural connection between the subiculum and the BNST in macaques and humans. 2) The spatial organisation of the structural connection between the subiculum and BNST in macaques and humans. 3) Whether individual differences in dMRI measures of white matter microstructure within subiculum - BNST tracts are phenotypically and/or genetically associated with self-reported stress-related traits. To do this, we first examined a series (n=7) of macaque monkeys with injections of tritiated amino acids in the hippocampal formation (Aggleton et al., 1986), to look for evidence of direct projections to the BNST. Secondly, we utilised an open-source macaque monkey dMRI dataset (n=9) to conduct in-vivo dMRI tractography analysis (Shen, Bezgin, et al., 2019a). Next, we repeated the dMRI tractography analysis in a large human sample, the Young Adults Human Connectome Project (HCP) (n = 1206). As well as containing high quality neuroimaging data (Sotiropoulos et al., 2013; Van Essen et al., 2012), this sample has been richly phenotyped; allowing for the examination of relationships between dMRI white matter microstructure and stress-related traits (reflecting dispositional negativity and alcohol use, derived in a previous study (Berry et al., 2020). Finally, we used the family structure of the HCP to assess the heritability, and co-heritability, of our dMRI microstructure measures and derived stress-related traits.

## 2. Methods

### 2.1 Data Descriptions

#### 2.1.1 Macaque Brain Tissue Data

For analysis of macaque neuroanatomical tract-tracing data we examined coronal sections from seven previously described adult cynomolgus monkeys (*Macaca fasicularis*) (Aggleton et al., 1986). The transport of tritiated amino acid from the hippocampal formation was visualised with autoradiography (Lanciego & Wouterlood, 2020). Although the tracer injections were made almost 40 years ago there is no evidence of a loss of signal, based on comparisons with drawings of label made shortly after initial tissue preparation. All experimental procedures were conducted consistent with the NIH Guide for Care and Use of Laboratory Animals (NIH Publication No. 86-23, revised 1985).

#### 2.1.2 Macaque Diffusion-MRI Data

Our dMRI monkey data were obtained from TheVirtualBrain Macaque MRI repository, made available through the OpenNEURO website (Shen, Bezgin, et al., 2019a). The sample consists of nine macaque monkeys - eight male rhesus macaques (*Macaca mulatta)* and one male cynomolgus macaque (*Macaca fasicularis)*. All surgical and experimental procedures were approved by the Animal Use Subcommittee of the University of Western Ontario Council on Animal Care and were in accordance with the Canadian Council of Animal Care guidelines (Shen, Bezgin, et al., 2019a).

#### 2.1.3 Human Diffusion-MRI Data

Human participants were drawn from the April 2018 release of the Young Adults Human Connectome Project study (n=1206) (Van Essen et al., 2012). Participants were between the ages of 25-37 and primarily made up of family groups, with an average size of 3-4 members and most containing a MZ (273) or DZ (166) twin pair. Participants were excluded during initial recruitment for psychiatric, neurological, or other long-term illnesses, although participants who were overweight, smoked, or had a history of recreational drug use and/or heavy drinking were included (Van Essen et al., 2012). For the imaging analysis, our samples included participants who had a dMRI and T1-w structural MRI scan (n=1065). Of these, there were 575 females and 490 males. For detailed recruitment information and for a full list of procedures see: https://www.humanconnectome.org/study/hcp-young-adult. See the supplementary material for a further breakdown of participants’ demographic information.

### 2.2 Hippocampal Injections

#### 2.2.1 Sample Preparation

Prior to the injection, the animals were sedated with ketamine hydrochloride (10 mg/kg), deeply anaesthetised with Nembutal (35 mg/kg) and placed within a stereotaxic apparatus. Under aseptic conditions, bone and dural flaps were opened to permit access to the temporal lobe. The animals received injections of an equal-parts mixture of tritiated proline (New England Nuclear, L-[2, 3, 4, 5 H], specific activity 139 Ci/mmole) and leucine (New England Nuclear L-[3, 4, 5 H], specific activity 111 Ci/mmole). Injections were made through a 1 µl Hamilton syringe at a final concentration of 50 µCi/µl. Following injection of the tracer, the dura and skin were sutured and the animals were allowed to recover. The analgesic, morphine (1 to 2 mg/kg subcutaneous every 4h) was given according to NIMH veterinary guidance. Recovery was without incident. Prophylactic doses of antibiotics were administered to prevent infection, and analgesics given during the post-operative period.

Single tracer injections of from 0.10 µl to 0.20 µl (5-10 μCi) were made in four cases (ACy12, ACy14, ACyF15, ACyF19), while two (ACy25, ACy28) of the remaining three monkeys received multiple injections within the same hemisphere totalling from 0.24 µl to 0.44 µl (12 μCi and 22 μCi). In one further case (ACyF27) injections were placed in both hemispheres. In the left hemisphere the injection was centred in the subiculum (ACyF27L), while the injection in the right hemisphere also involved the subiculum but extended into the caudal perirhinal cortex (ACyF27R). The coordinates for the hippocampal injection were determined with the aid of electrophysiological recordings made prior to the injection. A tungsten microelectrode was lowered into the hippocampal region and the various cell layers identified by their resting activity (Aggleton et al., 1986). Note that the hippocampal designations, including the subiculum, are based on the landmark work of Lorente de No (1934; see also Ding, 2013).

In three cases (ACyF15, ACyF19, ACyF27) the fornix was surgically sectioned months prior to the tracer injections. The surgical procedure and the completeness of the lesions have been fully documented elsewhere (Bachevalier et al., 1985).

After an interval of six - seven days the monkeys were sacrificed with a lethal dose of Nembutal (100mg/kg i.v.) and perfused through the left ventricle with normal saline followed by neutral buffered formalin. The brains were cryoprotected with 30% sucrose solution prior to being cut in 33 µm coronal sections on a freezing microtome. Every sixth section was mounted on a glass slide from phosphate buffer and then coated with Kodak NTB2 emulsion. The sections were exposed at 4°C for six to thirty weeks, developed in Kodak Dl9, fixed, and counterstained with thionine.

#### 2.2.2 Analysis of Subiculum to BNST Tract-tracing Data

Sections were examined under bright-field and dark-field using a Leica DM5000B microscope. Images of the regions of interest and the injection sites were acquired using Leica DFC310FX digital camera in the Leica Application Suite.

### 2.3 Diffusion-MRI Acquisition and Pre-processing

#### 2.3.1 Macaque Sample Image Acquisition

Full imaging acquisition protocols have been described elsewhere (Shen, Bezgin, et al., 2019a; Shen, Goulas, et al., 2019b). Briefly, the monkeys were anaesthetised before scanning and anaesthesia was maintained using 1-1.5% isoflurane during image acquisition. Images were acquired using a 7T Siemens MAGNETOM head scanner with an in-house designed and manufactured coil optimised for the Macaque head (Gilbert et al., 2016). Two diffusion-weighted scans were acquired for each animal, with each scan having opposite phase encoding in the superior-inferior direction at 1mm isotropic resolution. Six of the animals had data acquired with 2D echo planar imaging (EPI) diffusion and the remaining three had a multiband EPI diffusion sequence. Data were acquired for all animals with b=1000 s/mm^2^, 64 directions, 24 slices, and 2 b0 images (b-value = 0 s/mm^2^). A 3D T1w structural MRI scan was also collected for all animals (128 slices, 500 µm isotropic resolution) (Shen, Bezgin, et al., 2019a).

#### 2.3.2 Human (HCP) Diffusion-MRI Acquisition

All images were acquired on a 3T Skyra Siemens system using a 32-channel head coil, a customised SC72 gradient insert (100 mT/m) and a customised body transmit coil. High resolution anatomical images were acquired using a 0.7 mm isotropic T1-weighted 3D magnetisation-prepared rapid gradient echo sequence (TR 2,400 ms, TE 2.14 ms, FOV 224 × 224 mm^2^, flip angle 8°). Diffusion-weighted images were acquired using a spin-echo EPI sequence (voxel resolution 1.25 × 1.25 × 1.25 mm^3^, 111 slices, TR 5520 ms, TE 89.5 ms, flip angle 78°, refocusing flip angle 160°, FOV 210×180, echo spacing 0.78ms, BW 1488 Hz/ Px, b-values 1000, 2000, 3000 s/mm^3^ with 90 directions each and 18 b0 images (b-value = 0 s/mm^2^). Full details regarding acquisition parameters can be found in the HCP 1200 subject reference manual https://humanconnectome.org/study/hcp-young-adult/document/1200-subjects-data-release (Glasser et al., 2013; Sotiropoulos et al., 2013; Van Essen et al., 2012).

#### 2.3.3 Macaque Diffusion-MRI Pre-processing

The downloaded data had undergone pre-processing, including susceptibility-induced distortion correction using FSL’s topup and eddy tools (Andersson & Sotiropoulos, 2016; Jenkinson et al., 2012; Shen, Goulas, et al., 2019). Skull stripping was performed on the extracted B0 images using FSL BET (Jenkinson et al., 2012; Smith, 2002), BET parameters were optimised for each image (see supplementary materials). Binary masks of the extracted brains were multiplied with the respective 4D dMRI images to obtain skull-stripped diffusion weighted imaging (DWI) data (Jenkinson et al., 2012; Smith et al., 2004). All in-house scripts used for pre-processing with a step-by-step guide can be found online at https://github.com/El-Suri/Analyse-Monkey-Brain-Tracts.

#### 2.3.4 Human (HCP) Diffusion-MRI Pre-processing

The HCP data were downloaded having already undergone the standard minimal pre-processing pipeline for the Human Connectome Project (Glasser et al., 2013). Similar to the macaque imaging data, the subjects’ images were corrected for eddy currents and movement artifacts with the FSL eddy tool (Andersson & Sotiropoulos, 2016; Glasser et al., 2013). Images were skull-stripped using the FSL BET tool (Smith et al., 2004). The pipeline also included co-registration of subjects’ diffusion-weighted and anatomical scans. Full pre-processing steps and the code to run the HCP pre-processing pipeline can be found at https://github.com/Washington-University/HCPpipelines.

### 2.4 Region of Interest Selection

For both the macaque and human dMRI data we extracted several regions of interest (ROIs) for our analyses. The seed region for all analyses was the subiculum ROI. The subiculum ROI here represents the subiculum complex, a term that incorporates the prosubiculum, subiculum, the presubiculum, and the parasubiculum (de Nó, 1934; Ding, 2013; O’Mara et al., 2001). The principal target region of interest was the BNST. We had two positive comparison target regions, the anterior thalamic nuclei (ATN) and nucleus accumbens (NAc), both of which are located in close proximity to the BNST and show substantial structural connectivity with the subiculum, as revealed by tract-tracing (Aggleton & Christiansen, 2015; Christiansen et al., 2016; Friedman et al., 2002). For a negative comparison region we selected the external globus pallidus (GPe), as this is also within close proximity of the BNST but does not appear to directly connect with the subiculum (Aggleton & Christiansen, 2015).

Some findings suggest that the primate fornix is not necessary for all subiculum to BNST connectivity, with evidence implicating a secondary path via the amygdala, involving the stria terminalis and/or the amygdalofugal pathway (Cullinan et al., 1993; Poletti et al., 1973, 1984; Poletti & Creswell, 1977; Poletti & Sujatanond, 1980). To assess this possibility, we ran an additional analysis using a fornix mask to exclude any connections passing through this region.

Although we expected some deviation in the definitions and boundaries of the ROIs between the macaque and human atlases, we tried to ensure that the structures analysed were closely comparable and that the relative sizes of the structures in the two species were as similar as possible. See Figures 1 & 2 for a visual comparison of the ROIs, which are described further below.

**Figure 1.**
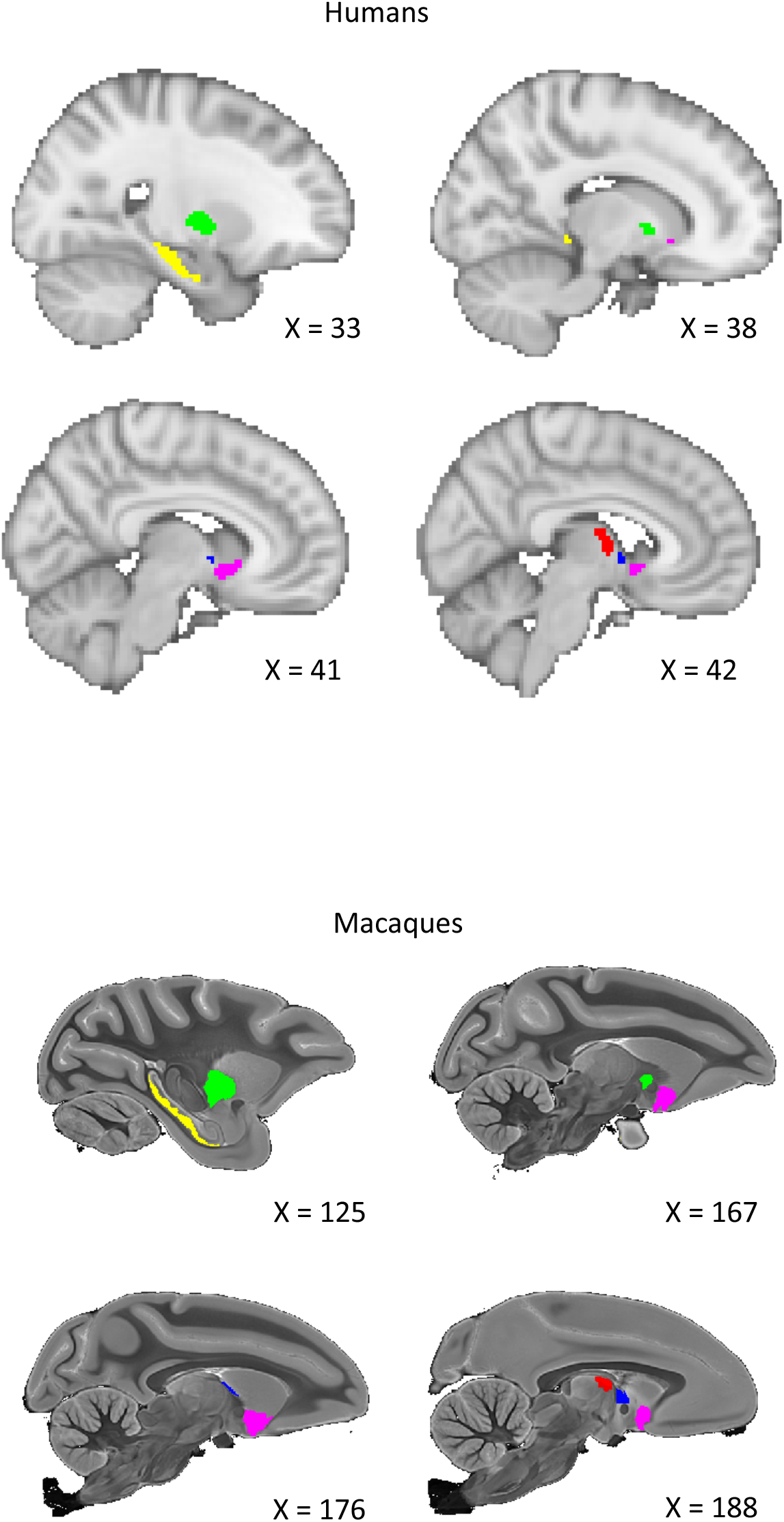
Sagittal view of the regions of interest for humans and macaques. Yellow = Subiculum, Green = External Globus Pallidus (GPe). Red = Anterior Thalamic Nuclei (ATN), Blue = BNST, Purple = Nucleus Accumbens (NAc). Coordinates for the macaque are taken from the atlas of Calabrese et al. (2015).

**Figure 2.**
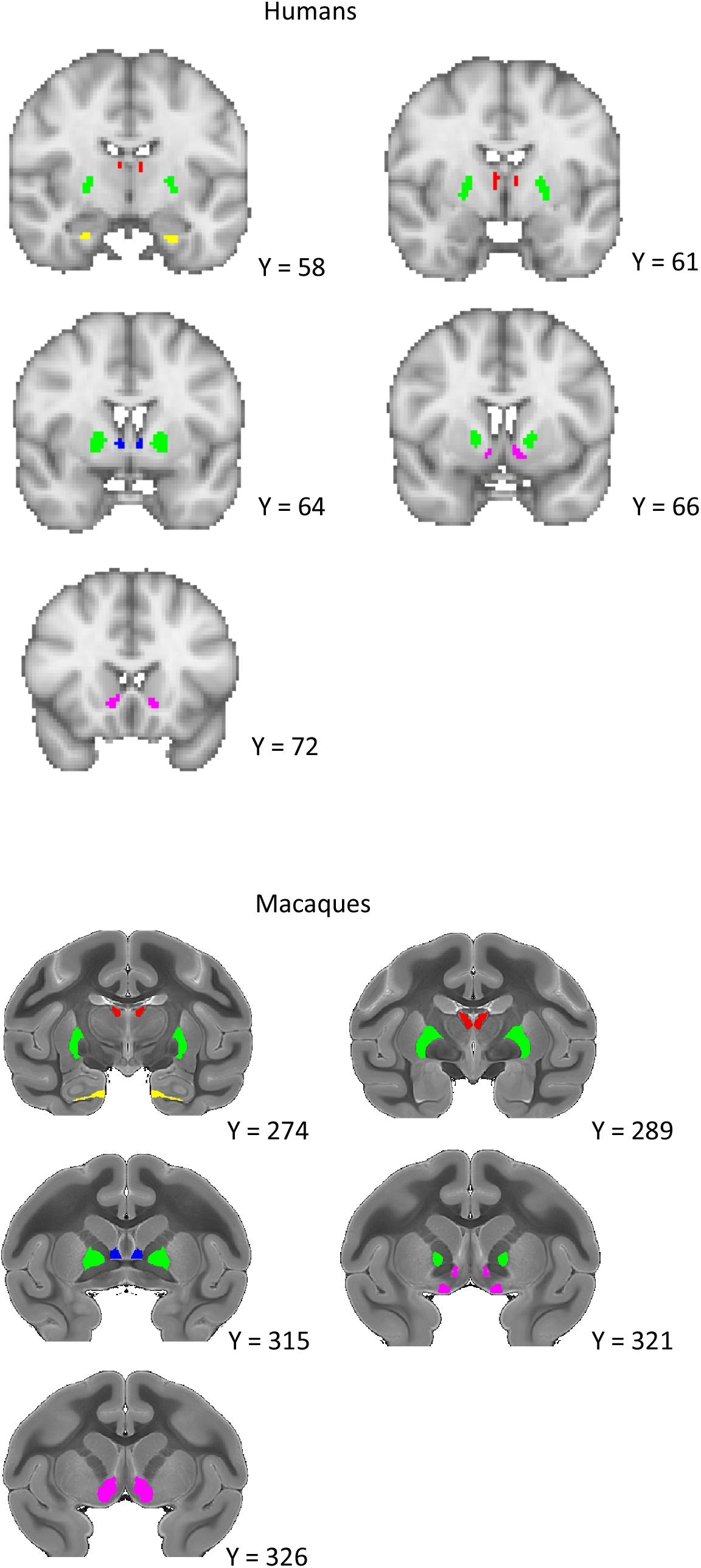
Coronal view of the regions of interest for humans and macaques. Yellow = Subiculum, Green = External Globus Pallidus (GPe). Red = Anterior Thalamic Nuclei (ATN), Blue = BNST, Purple = Nucleus Accumbens (NAc). Coordinates for the macaque are taken from the atlas of Calabrese et al. (2015).

#### 2.4.1 Macaque Anatomical Regions of Interest

To create subject-specific brain ROIs we registered a standardised macaque brain atlas to individual subjects’ native diffusion space. We used a high-resolution rhesus macaque MRI atlas containing 241 brain regions, created from the imaging of 10 post-mortem specimens (Calabrese et al., 2015). Although we had T1-w anatomical images for our macaque sample, the atlas included only T2-w anatomical images, and so to register the atlas to the individual macaque space we used the B0 images from both datasets. Registration was carried out using the FSL FLIRT tool (Jenkinson et al., 2012; Smith et al., 2004). To check the results of this registration, we registered the macaque T1-w scans to their respective B0 images and overlayed the T1-w image to the now subject-space atlas label images for visual inspection. An in-house script was then used to extract the specific brain ROIs from the subject-space atlases. All regions were identified from a published atlas (Calabrese et al., 2015). As previously mentioned, for the subiculum ROI we combined the prosubiculum, subiculum, presubiculum, and parasubiculum ROIs into a single mask (de Nó, 1934; Ding, 2013; O’Mara et al., 2001). For the NAc, the NAc core and NAc shell ROIs were combined into a single mask. The ATN mask was a combination of the anteroventral and anteromedial thalamic ROIs. These two regions have been reported to have strong connectivity with the subiculum and together form a structure similar in size to the BNST (Aggleton & Christiansen, 2015). For the BNST, we used the central BNST ROI but manually removed the portion below the anterior commissure within FSL. This was done so that it matched our human BNST mask, which only contains the dorsal BNST (Theiss et al., 2017; Torrisi et al., 2015).

#### 2.4.2 Human Anatomical Regions of Interest

All HCP ROIs were defined in 2mm MNI space and then transformed to the subject diffusion space during the FSL PROBTRACKX2 analysis (see section 2.5.2) via the non-linear transformations provided by the HCP (Behrens et al., 2007). The subiculum and NAc ROIs were taken from the Harvard Oxford Subcortical atlas and, to better match the macaque ROIs, were thresholded to a minimum probability of 80% and 50% respectively (Jenkinson et al., 2012; Smith et al., 2004). The subiculum mask provided by the Harvard Oxford Subcortical atlas does not distinguish between the subicular sub-regions, and thus ‘subiculum ROI’ also refers to the larger subiculum complex (de Nó, 1934; Ding, 2013; O’Mara et al., 2001). The BNST mask was downloaded from the NeuroVault website (https://neurovault.org/collections/3245), originally uploaded by Theiss et al (Mai et al., 2015; Theiss et al., 2017; Tillman et al., 2018). This mask was thresholded to a minimum probability of 25%, in accordance with previous studies (Berry et al., 2020; Theiss et al., 2017). The thalamic nuclei mask centred on the anteroventral ROI taken from the Automated Anatomical Labelling Atlas 3 (Rolls et al., 2020). This ROI includes the anteromedial area, thus matching the NHP sample (Iglesias et al., 2018; Rolls et al., 2020). The GPe ROI was downloaded from the NeuroVault website (https://identifiers.org/neurovault.collection:1380) and transformed to MNI 2mm space using FSL FLIRT (Jenkinson et al., 2012; Smittenaar et al., 2017). To better match the macaque ROI, this mask was thresholded at 50%. For our fornix mask, we used the fornix column and body mask from the ICBM-DTI-81 white-matter labels atlas in FSL (Hua et al., 2008; Jenkinson et al., 2012; Mori et al., 2008; Wakana et al., 2007). See Figures 1 & 2 for a comparison of the macaque and human seed and target ROIs.

### 2.5 Probabilistic Diffusion Tractography

#### 2.5.1 Tractography Analyses

White matter tractography was performed using FSLs (v 6.0.1) BEDPOSTX and PROBTRACKX2 (Behrens et al., 2007; Jenkinson et al., 2012). Briefly, BEDPOSTX uses Markov Chain Monte Carlo sampling to build up distributions on diffusion parameters. A 2-fibre model was applied to improve the modelling and resolution of crossing fibre populations (Behrens et al., 2007). PROBTRACKX2 then uses the outputs of BEDPOSTX to model the probabilities of white matter pathways in the brain. It does this by repeatedly initiating streamlines from a given seed region, iterating between drawing an orientation from the BEDPOSTX distributions, taking a step in this direction, and checking for any termination criteria (Behrens et al., 2007). The streamline distributions can be used to estimate the probability of structural connectivity between brain regions (see section 2.5.2) and image potential pathways (computational tract-tracing). For both our macaque and human samples we ran an ROI x ROI analysis. Although we were only interested in the subiculum ROI as the seed region, this analysis allowed us to run all subiculum-target ROI combinations at once. For the HCP sample only, to assess the contribution of the fornix to subiculum ROI - BNST connectivity, we ran two subiculum ROI - BNST analyses, one of which used a fornix mask as a NOT gate. This meant that any streamlines passing into the fornix would be eliminated from this analysis. Because our target ROIs varied in their distance from our seed, and because the distance between ROIs is a factor influencing successful reconstruction of a connection, we applied the distance correction setting within PROBTRACKX2 to account for this (Behrens et al., 2007). Probabilistic fibre tracking was initiated from every voxel within the ROI mask, with 5000 streamline samples being sent out from each voxel. We applied a step length of 0.2mm (NHP) or 0.5mm (HCP), and a curvature threshold of 0.2. For the full list of the parameters used during the BEDPOSTX and PROBTRACKX2 analyses see the supplementary materials.

#### 2.5.2 PROBTRACKX2 Data Transformation

The outputs of the PROBTRACKX2 analysis consist of an ROI by ROI connectivity matrix. Each number in the connectivity matrix refers to the total number of streamlines that were successfully reconstructed between each pair of ROIs. In this analysis, each ROI is treated as both a seed and a target, meaning that a reconstruction is run twice for every ROI combination. The mean of each ROI to ROI combination was taken for a more robust estimate of the connectivity probability (Gschwind et al., 2012). As we were only interested in connectivity between the subiculum and the other ROIs, we did not use the data regarding other ROI to ROI streamlines (e.g., BNST to NAc). In order to express each subiculum to target ROI connectivity value as a proportion, relative to each subjects total number of subiculum streamlines, each mean subiculum - target ROI value was divided by the total number of streamlines successfully propagated from the subiculum for each participant (Gschwind et al., 2012). This gave us a proportion of connectivity for each seed-target combination between 0 and 1 (Gschwind et al., 2012). Therefore, we refer to this measure as a connection proportion.

#### 2.5.3 Statistical Analysis of Connection Proportion Differences

To test whether the connection proportion between the seed and target regions were different, we ran two within-groups one-way ANOVA’s, one for each species, with Target ROI as the IV and Connection Proportion as the DV. Outliers were removed if they were more than 1.5 * the inter-quartile range (IQR) above the third quartile or below the first quartile (e.g., Vinutha et al., 2018). Mauchly’s test of sphericity and Levene’s test of equality of error variances were run. Subsidiary pairwise t-tests were performed to assess the nature of the ROI differences. P-values for all subsidiary tests were corrected via the Bonferroni method.

#### 2.5.4 Statistical Analysis of Between-Species Differences

To statistically test whether the subiculum ROI connectivity patterns of the NHP and human samples were different, we ran a *post-hoc* 2×4 mixed ANOVA, with Species as the between-groups variable and Target ROI as the within-groups variable.

#### 2.5.5 Statistical Analysis of Fornix Exclusion Effects on Subiculum to BNST Connectivity

Finally, we compared the number of streamlines that were successfully reconstructed between the subiculum ROI and BNST when a route via the fornix was, versus was not, available. For this comparison, we used the raw number of streamlines created between the two ROIs (i.e., not the connectivity proportion) when a) our fornix exclusion mask was not applied, and b) when our fornix exclusion mask was applied. These two sets of values were compared using a paired-samples t-test.

### 2.6 White Matter Microstructure and Stress-Related Traits and Behaviours

#### 2.6.1 Principal Component Analysis of Stress-related Traits and Behaviours

To assess the association between measures of white matter microstructure and stress-related traits and behaviours, we utilised the HCP self-report questionnaire data relating to these constructs (Weintraub et al., 2013). In previously published work we used principal component analysis (PCA) to reduce nine questionnaire items relating to anxiety, depression, perceived stress, fear, and substance use to two principal components, one representing ‘Dispositional negativity’ and one ‘Alcohol use’ (see Berry et al., 2020 for details). These two principal components were thus used as our phenotypes of interest.

#### 2.6.2 White Matter Microstructure

For our measures of white matter microstructure we utilised the tensor model, which assesses the properties of water diffusion along a tensor to infer the microstructural properties of white matter pathways (Basser, 1997). We extracted Fractional Anisotropy (FA), Mean Diffusivity (MD), Axial Diffusivity (AD), and Radial Diffusivity (RD) from the subiculum to BNST tracts. To do this we ran FSLs DTIFIT command on each subject’s DWI data, providing FA and MD, and AD values across the brain. RD values were calculated by adding the L2 to the L3 image and dividing by two (Winklewski et al., 2018). A new PROBTRACKX2 analysis was then run with the subiculum ROI as the seed and the BNST as the only target. The resulting brain image, containing the probability distribution of connections between the two regions, was thresholded so that voxels with a less than 10% probability were excluded. This image was then binarized and transformed to subject space, where it was used to extract the mean FA, MD, AD, and RD values of the tracts.

#### 2.6.3 Heritability and Association Analyses

We used the SOLARIUS package for R to assess the following: 1) the heritability of white-matter microstructure (FA, MD, AD, and RD) within the subiculum ROI - BNST tracts, 2) the co-heritability of these metrics with the dispositional negativity and alcohol use components, 3) the phenotypic, genetic, and environmental correlations between the subiculum-BNST white matter microstructure and component scores for dispositional negativity and alcohol use. SOLARIUS is the R version of the widely used SOLAR-eclipse software for genetic analysis (O’Donnell & Westin, 2011). SOLAR uses a kinship matrix to estimate the proportion of variance in a phenotype attributable to additive genetics, the environment, or to residual error. In this case, we were only permitted to calculate the additive genetic component, as partitioning environmental and error effects requires household information that is not provided by the HCP. In this model, monozygotic twins are given a score of 1 and dizygotic twins / siblings of 0.5 to indicate the estimated proportion of shared genetic variation. Half-siblings were excluded from the analysis (n = 191). The pedigree file was created using the HCP2Solar MATLAB function, a tool specifically designed for the HCP participants (https://brainder.org/2016/08/01/three-hcp-utilities) (Winkler et al., 2015). Because the model is sensitive to kurtosis, the phenotype values were inverse normally transformed. SOLARIUS allows analysis of co-heritability by computing bi-variate genetic correlations (Kochunov et al., 2019). During the analysis SOLARIUS computes an estimate of phenotypic, genetic, and environmental correlation between the variables, which we used to assess the relationships between white matter microstructure and the dispositional negativity and alcohol use components. The covariates for all analyses were sex, age, BMI, age^2^, sex * age, and sex * age^2^. The final number of participants in these analyses was n = 933. For discussion on using SOLAR for genetic neuroimaging see (Kochunov et al., 2019).

## 3. Results

### 3.1 Macaque Subiculum – BNST Neuroanatomical Tract-tracing

#### 3.1.1 Evidence for Direct Projections from Autoradiography

Information came from seven cases with injections of amino acids within the hippocampal formation. In all cases, except for ACyF19 and ACy25, the injection sites involved the subiculum. The clearest evidence of a direct projection to the BNST was seen in ACy28, where an extensive injection filled, but remained restricted, within the caudal hippocampal formation. Consequently, the injection included the caudal CA fields, dentate gyrus, presubiculum, and subiculum, reaching the proximal border of the presubiculum (Figure 3).

**Figure 3.**
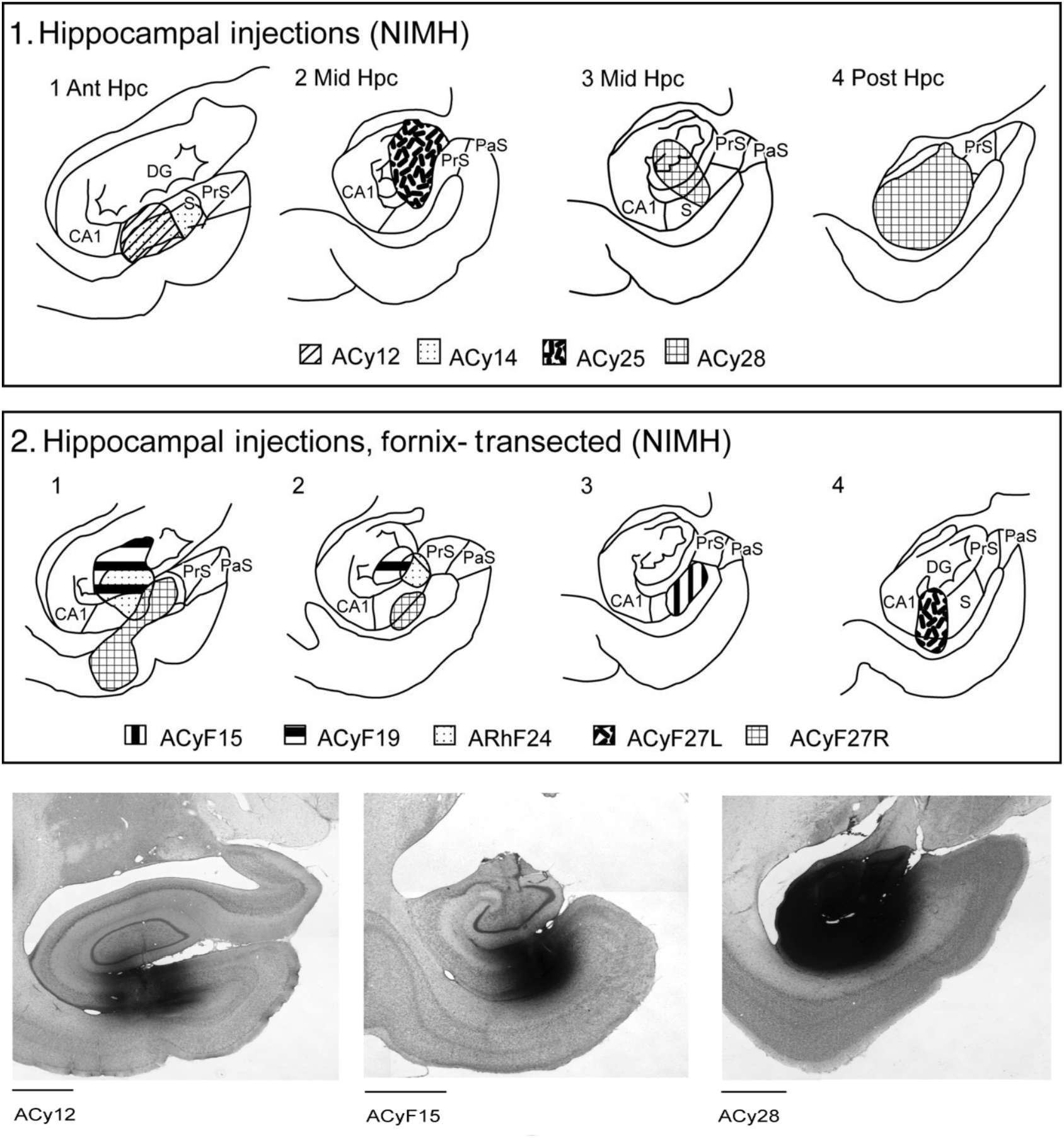
The upper two rows depict the core of each amino acid injection in the hippocampal formation drawn onto standard coronal sections at different anterior-posterior levels (all *Macaca fascicularis* cases). Row 1, intact cases. Row 2, cases with prior fornix transection. Row 3 contains coronal photomicrographs of the centre of the amino acid injection in three cases, at anterior, mid, and posterior levels, respectively. CA1, hippocampal field CA1; DG, dentate gyrus; Hpc, hippocampus; NIMH, National Institute of Mental Health; PaS, parasubiculum; PrS, presubiculum; S, subiculum. Scale bar = 2mm

In ACy28 fornical inputs to the BNST could be readily seen (Figure 4). A very visible patch of localised label (including fibres) was present dorsal to the rostral part of the anterior commissure (Figure 4). This region corresponds to the medial division of the anterior part of the BNST (Paxinos et al., 2009). Only limited BNST label was seen ventral to the anterior commissure, much of which comprised fibres. Some fibres appeared to pass through and around the more caudal BNST to reach hypothalamic targets. No labelled fibres could be seen in the stria terminalis itself.

**Figure 4.**
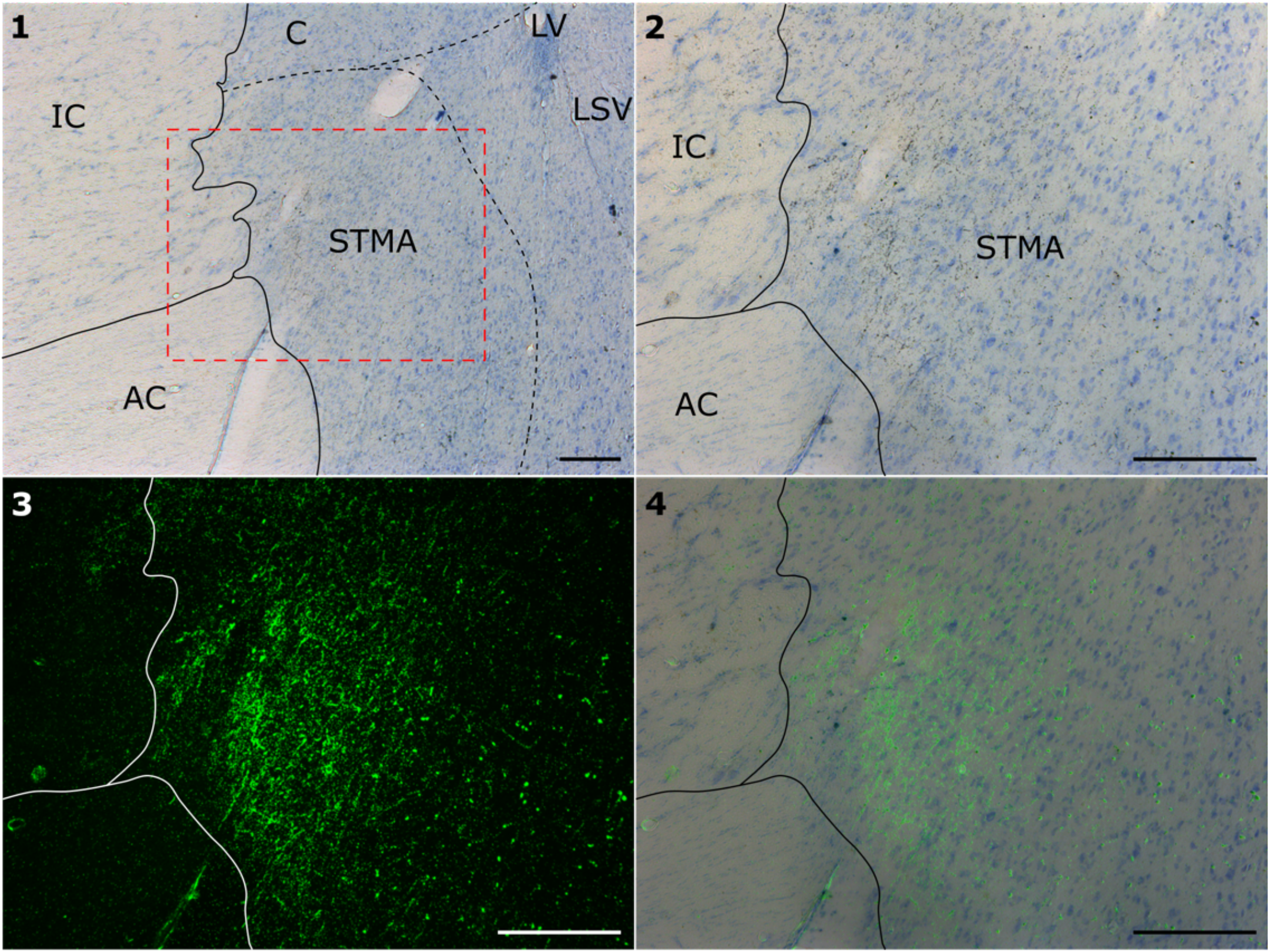
Bright field (#1, #2), dark field (#3), and overlay (#4) showing anterogradely transported label (green) in the bed nucleus of the stria terminalis (BNST) following tracer injections in the posterior hippocampal formation of case ACy28. Image #2 is within that part of image #1 outlined with red dashes. Abbreviations: AC, anterior commissure; C, caudate nucleus; IC, internal capsule; LSV, lateral septum, ventral; LV, lateral ventricle; STMA, bed nucleus of the stria terminalis, medial division, anterior. All scale bars 250µm.

Two cases had tracer injections centred in the rostral subiculum and prosubiculum (ACy12, ACy14). In both cases, dense fornical label could be followed to the level of the anterior commissure. Labelled fibres could then be observed following through and immediately around the BNST, accompanied by light, diffuse label consistent with termination in the medial division of the anterior part of the BNST. This label was especially light in ACy14. Again, no label could be seen in the stria terminalis. Little or no label was detectable ventral to the anterior commissure within the BNST in these two cases. Finally, there was no evidence of any BNST label in case ACy25, in which the injection involved the dentate gyrus and CA3, but not the subiculum. Consequently, the BNST label was associated with involvement of the subiculum within the injection site.

There was no evidence of label in the BNST in the three cases with fornix transection (ACyF15, ACyF19, ACyF27L). Of these three cases, terminal label could be seen in the basal amygdala in both ACyF15 and ACyF27 (see Aggleton, 1986). In none of the seven cases was label apparent in the stria terminalis.

### 3.2 Subiculum Complex – BNST Probabilistic dMRI

3.2.1 Macaque Tractography Results

Macaque tractography analysis seeded from the subiculum ROI demonstrated the highest proportion of connectivity to the ATN, followed by the BNST, the NAc, and the GPe (Figure 5). ANOVA revealed a significant effect of ROI (F(3,12) = 29.42, p <0.0001, Nagelkerke r^2^ = 0.88), with follow-up t-tests demonstrating a statistically significant difference between the ATN and the other regions (Table 1). No other subiculum-target ROI combinations were significantly different from each other (Table 1). Viewing the tractography output images, after thresholding by 10%, most reconstructed tracts appeared within the fornix, though some also passed through the amygdala via an area consistent with the amygdalofugal pathway (Figure 6). There was no evidence for connectivity beyond the anterior commissure at this threshold. An interactive combined output image, consisting of all 9 NHP tractography results, has been uploaded to NeuroVault at https://identifiers.org/neurovault.image:682610.

**Table 1.**
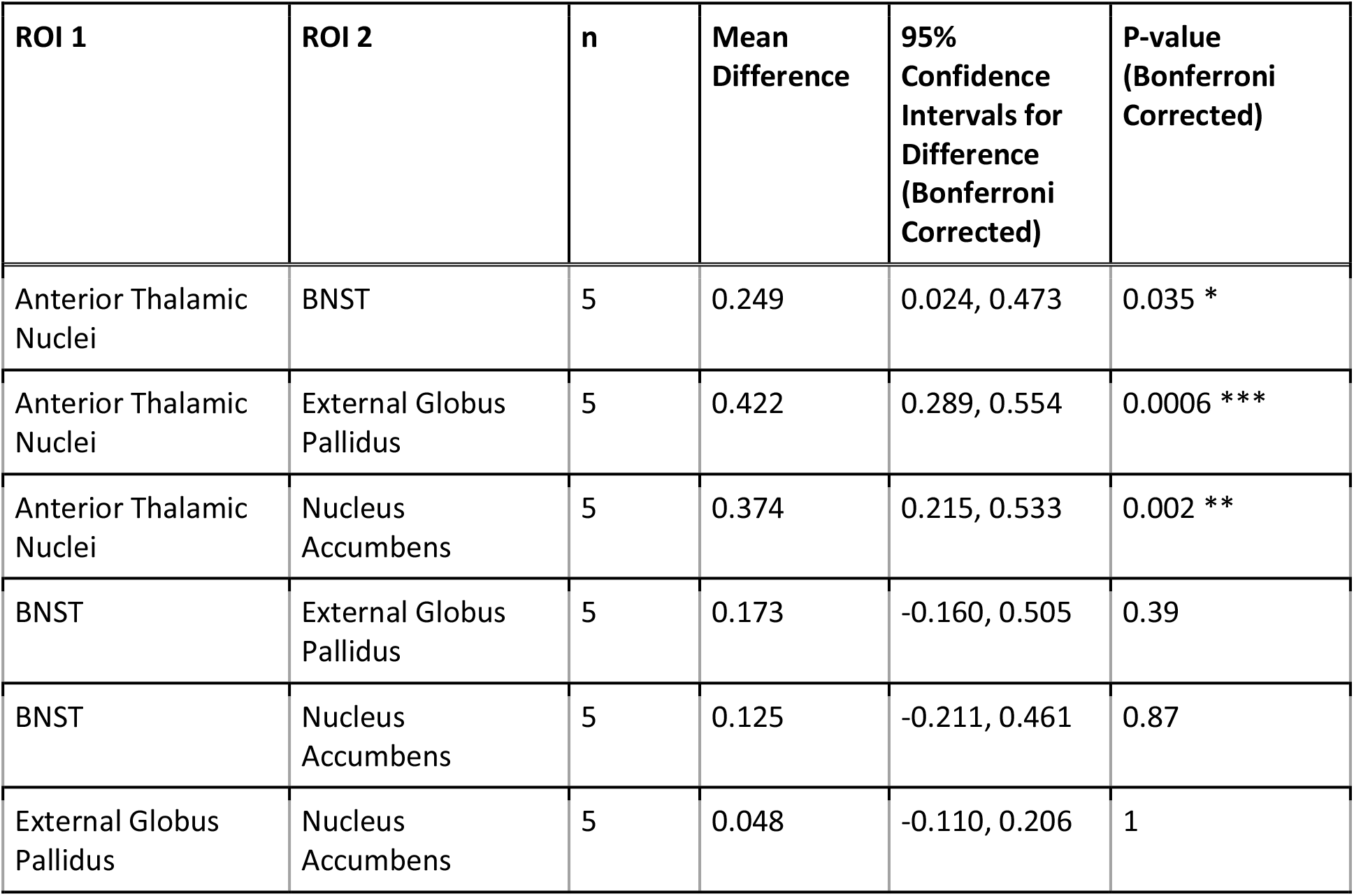
Macaque (top) and human (bottom) t-test demonstrating the differences between the connection proportion of each subiculum – region of interest (ROI) pair. BNST = Bed Nucleus of the Stria Terminalis.

**Figure 5.**
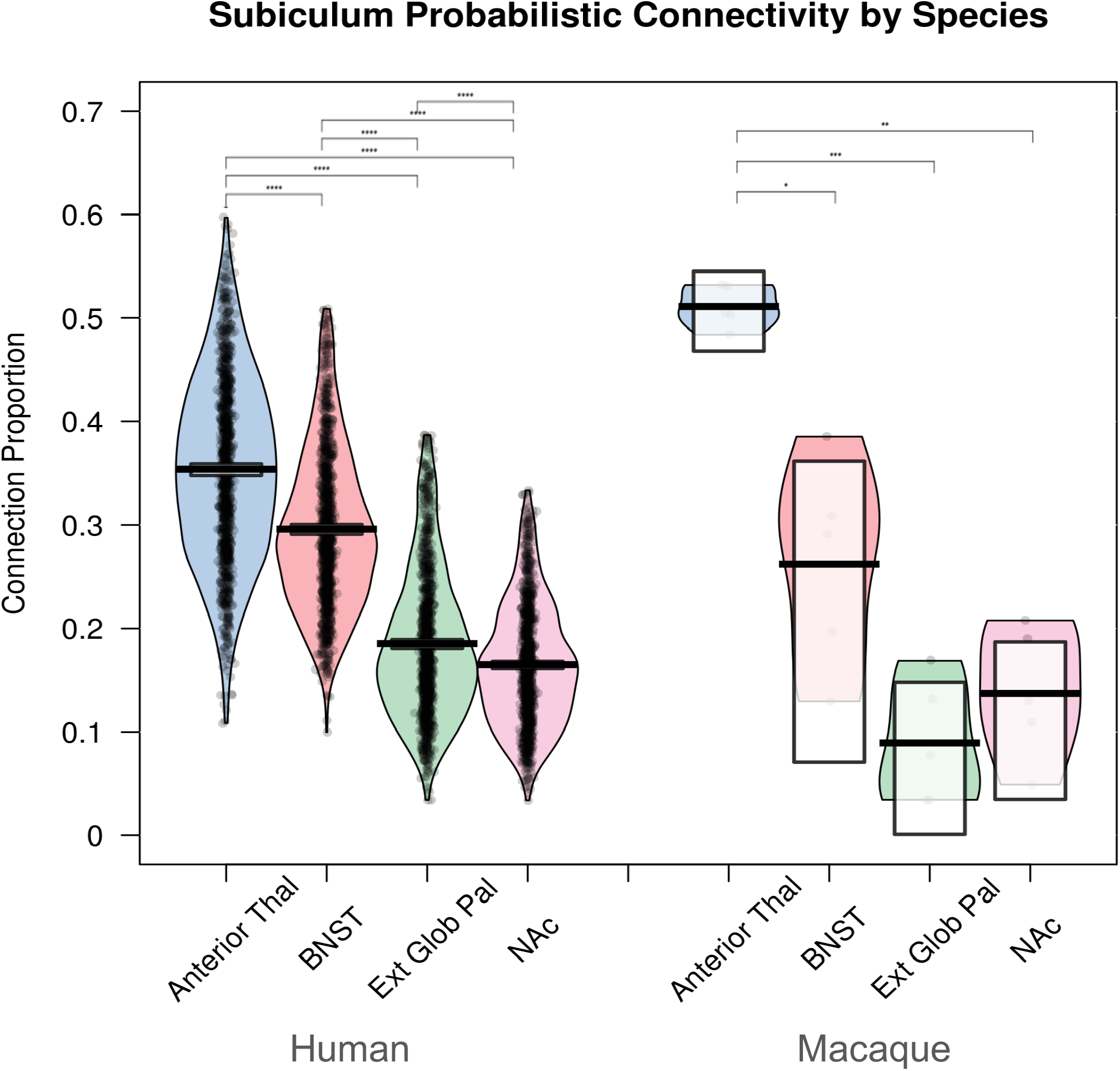
The proportion of connectivity from the subiculum with each of the target regions for macaques and humans. Broadly the same pattern is seen across species, with the highest proportion of connectivity with the anterior thalamic nuclei (ATN), followed by the BNST, and then either the external globus pallidus (GPe) or nucleus accumbens (NAc). A mixed ANOVA revealed that there was a significant interaction of Species x ROI, but no main effect of Species. These ROI differences were highly significant after Bonferroni correction for every ROI in the HCP sample (n=985), but only for the anterior thalamic nuclei (ATN) in the NHP sample. (n=5).

**Figure 6.**
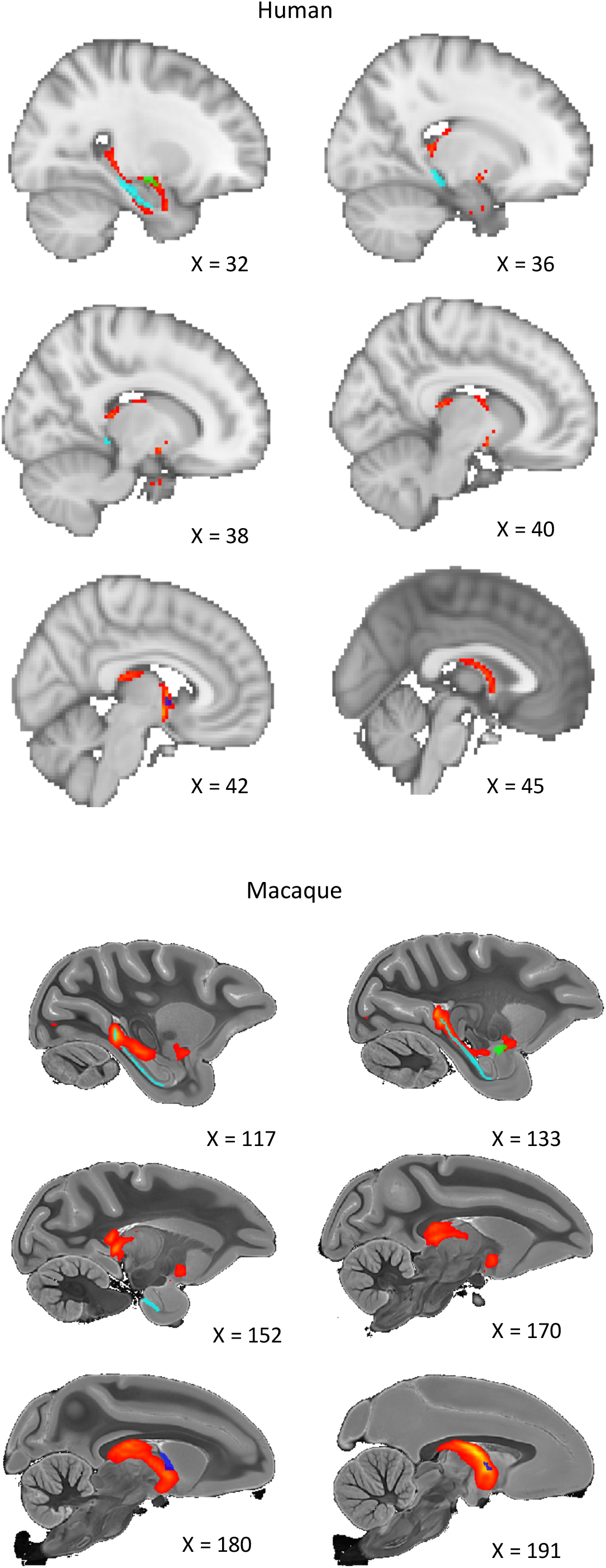
Human and macaque sagittal images from the probabilistic tractography analysis, seeded from the subiculum (light blue) with the BNST (dark blue) as the target. The connection is primarily via the fornix, however, significant connectivity is also observed through the amygdala (green). Images are thresholded at 10%.

#### 3.2.2 Human Tractography Results

Human dMRI tractography analysis seeded from the subiculum ROI demonstrated the highest proportion of connectivity to the ATN, followed by the BNST, the NAc, and the GPe (Figure 5). ANOVA revealed a significant effect of ROI (F(2.65, 2612.46) = 1069.56, p < 0.0001, Nagelkerke r^2^ = 0.52), with subsidiary pairwise comparisons demonstrating significant differences between all subiculum-target combinations (Table 2). Viewing the 10% thresholded tractography output image, the tractography findings were very similar to that of the macaques, with most of the tracts passing via the fornix but not extending in advance of the anterior commissure (Figure 7). An area of connectivity was also seen through the amygdala, via a route consistent with the amygdalofugal pathway. An interactive combined output image, consisting of the first 10 HCP tractography results, has been uploaded to NeuroVault at https://identifiers.org/neurovault.image:682612.

**Table 2.**
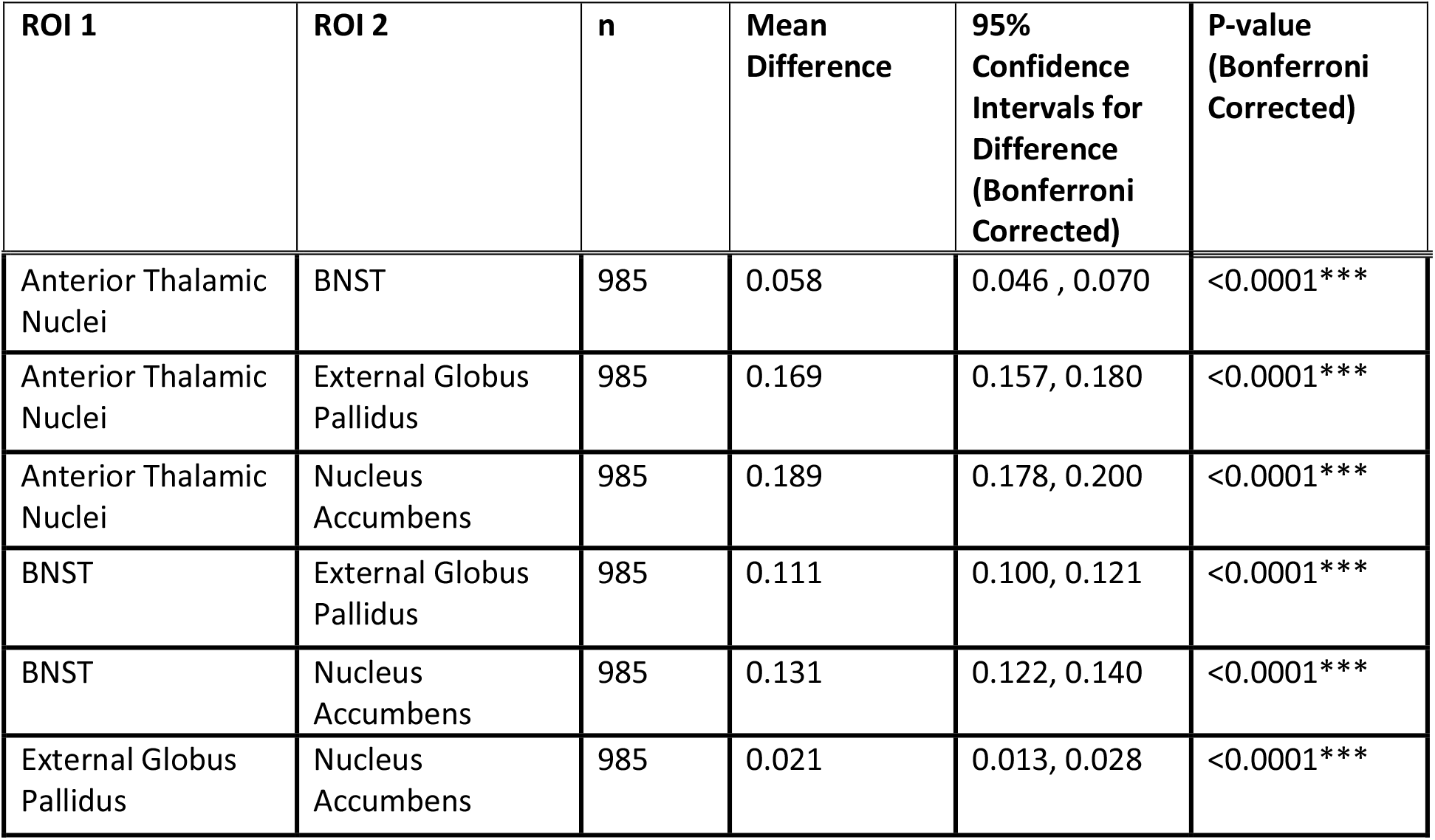
Macaque (top) and human (bottom) t-test demonstrating the differences between the connection proportion of each subiculum – region of interest (ROI) pair. BNST = Bed Nucleus of the Stria Terminalis.

**Figure 7.**
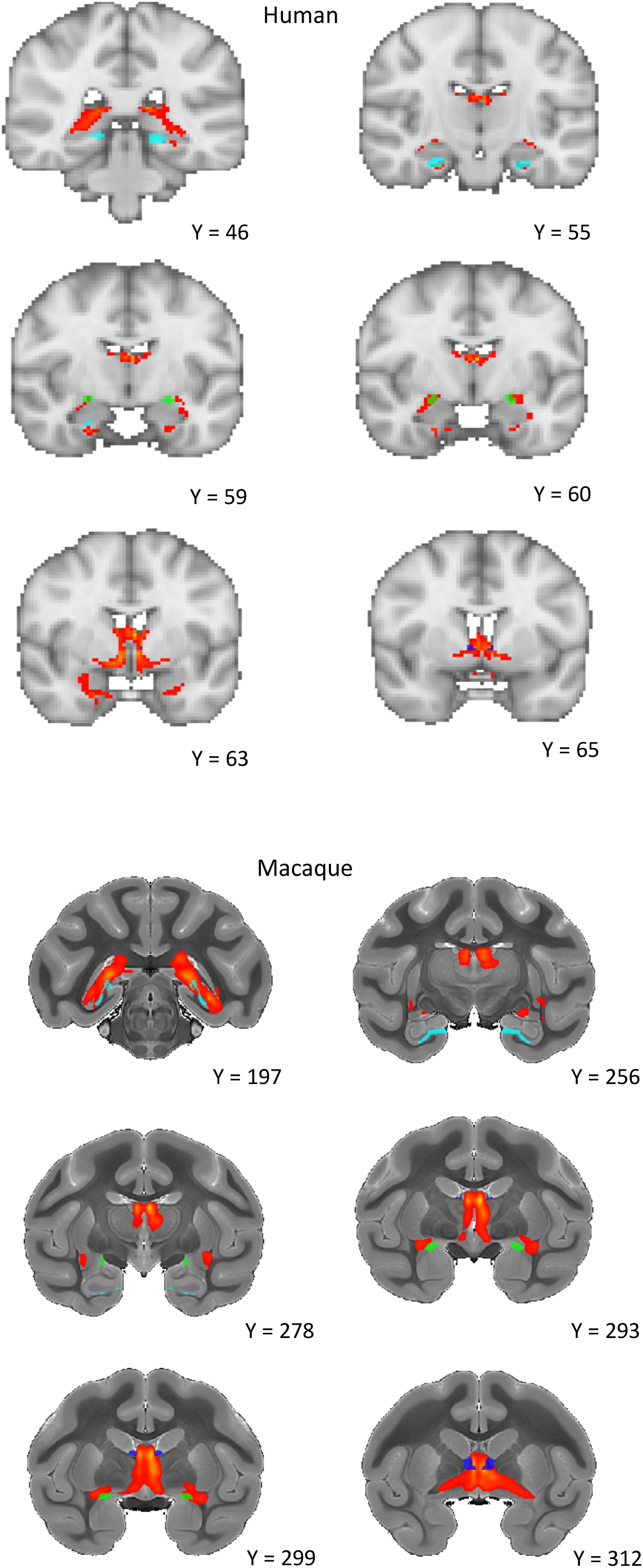
Human and macaque coronal images from the probabilistic tractography analysis, seeded from the subiculum (light blue) with the BNST (dark blue) as the target. The connection is primarily via the fornix, however, significant connectivity is also observed through the amygdala (green). Images are thresholded at 10%.

#### 3.2.3 Statistical Comparison of Macaque and Human Subiculum Complex Tractography Results

The macaque and human dMRI tractography results showed largely the same pattern of connectivity, with the highest proportion of streamlines linking the subiculum ROI to the ATN, followed by the BNST, and then either the NAc or GPe (although statistically significantly more to the GPe in the HCP data) (Figure 5). Statistical analysis to formally assess these differences found a significant interaction of Species x ROI (F(2.66, 2626) = 7.95, p < 0.0001, ηp2 = 0.008, Greenhouse-Geisser corrected), with a significant main effect of ROI (F(2.66, 2626) = 50.3, p < 0.0001, ηp2 = 0.048, greenhouse-Geisser corrected) but no significant main effect of Species (F(1, 988) < 0.001, p = 1, ηp2 = 0). This indicates that there is a difference between the species regarding which ROI x ROI combinations were significantly different. The direction of these effects is described in the previous sections (also see Figure 5).

#### 3.2.4 Fornix Exclusion Results In Significantly Fewer Streamlines

Tractography of the subiculum ROI to the BNST that included the fornix contained significantly more streamlines (M = 5005, SD = 2827) than tractography without the fornix (M = 3348, SD = 2030) (t = 38.4, df = 988, p < 0.0001, Cohen’s d = 1.2) (Figure 8).

**Figure 8.**
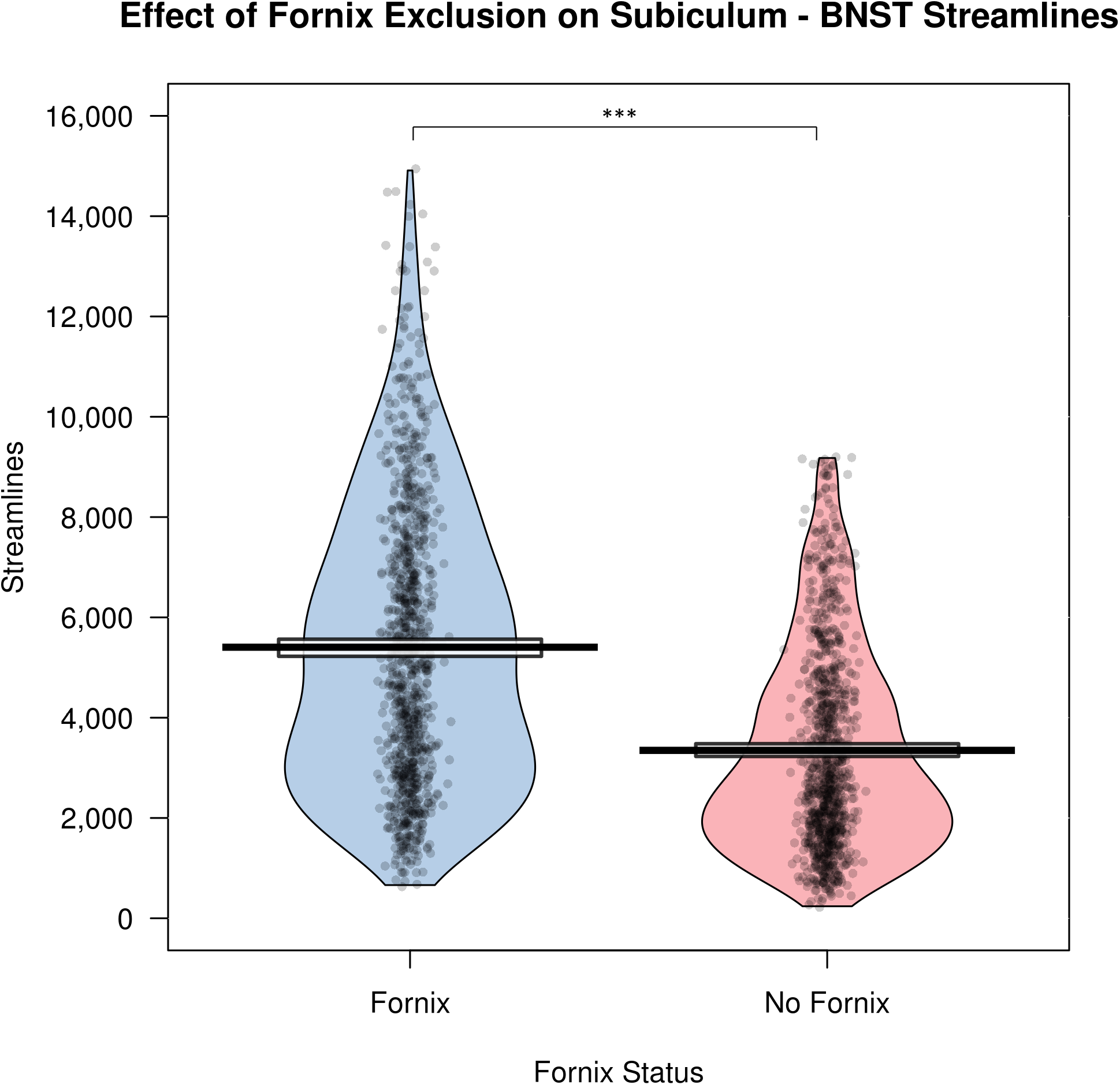
Excluding the fornix from the tractography analysis between the subiculum and BNST in the HCP sample made a significant difference to the number of streamlines between the two regions. Interestingly, despite the removal of this major subiculum output pathway, there remains substantial connectivity with the BNST.

### 3.3 Subiculum Complex - BNST Tract White Matter Microstructure Heritability and Phenotypic Associations

#### 3.3.1 Univariate Heritability Analysis

SOLAR analysis revealed that tensor-derived microstructure indices (FA, MD, RD, and AD) extracted from the subiculum ROI - BNST tracts were all significantly heritable (p<0.0001 h^2^ ∼.50) (Table 3). This was also the case for the subiculum to BNST connections excluding the fornix (P<0.0001 h^2^ ∼ 0.5) (Table 3).

**Table 3.**
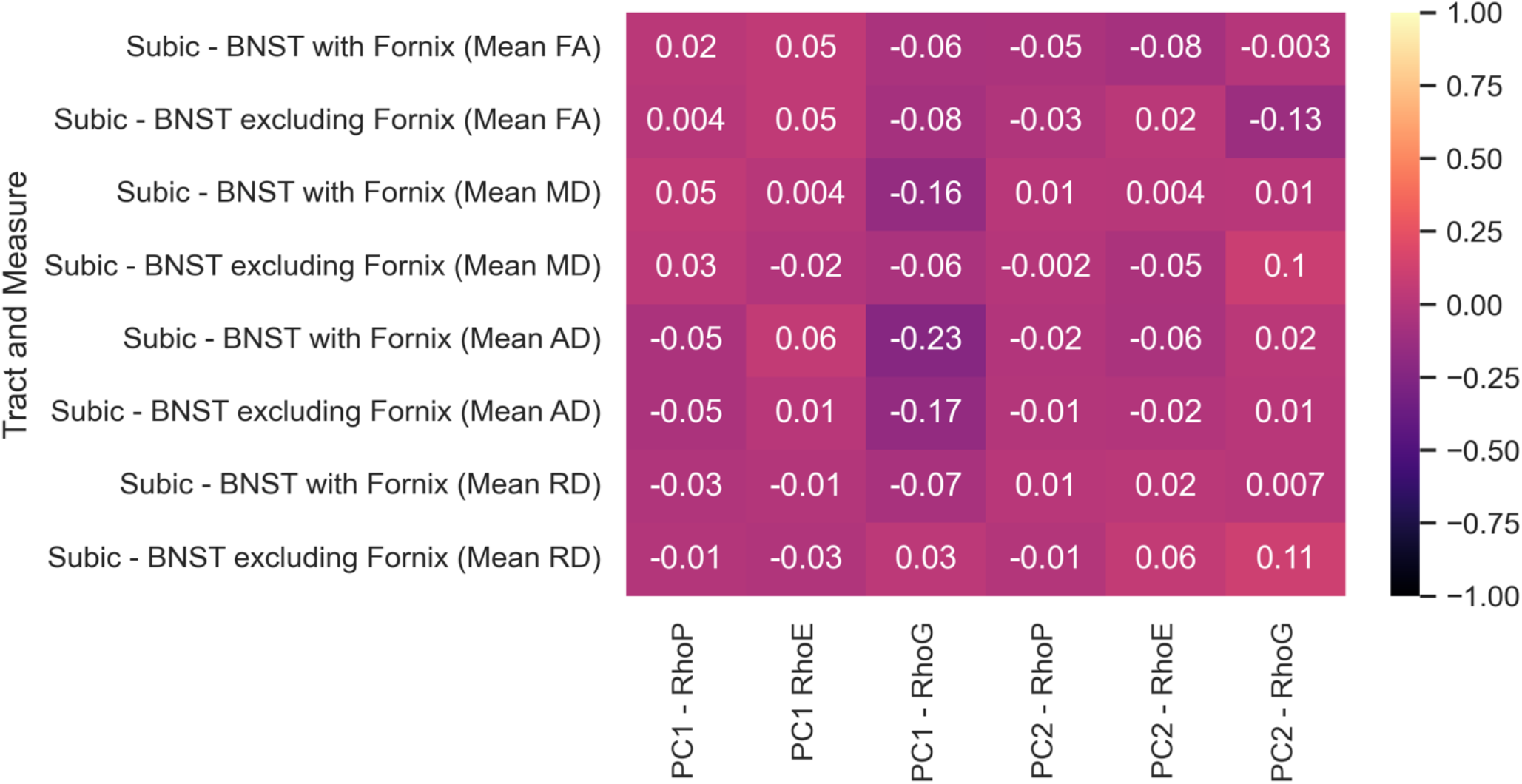
Correlation heatmap demonstrating the phenotypic (RhoP), environmental (RhoE), and genetic (RhoG) correlations following bivariate heritability analyses between each extracted DTI measure and each principal component (PC). The analysis revealed no significant correlations between any of the measures (p > 0.05). PC1 represented traits related to dispositional negativity, whereas PC2 reflected measures of alcohol use. See supplementary methods for p-values and standard errors.

#### 3.3.2 Bivariate Heritability and Association Analysis

Bivariate heritability analysis for each microstructure measure with each of the two behavioural components (dispositional negativity and alcohol use) revealed no evidence for phenotypic, environmental, or genetic associations between any of the variables (p > 0.05, Table 4). This was also the case for the subiculum ROI - BNST connections excluding the fornix (p > 0.05, Table 4). However, the covariates BMI, age, and sex were highly significantly associated with the microstructural measures, within tracts both including and excluding the fornix (Table 4).

**Table 4.** Univariate heritability analysis demonstrated that each DTI microstructure measure extracted from the subiculum to BNST connection was moderately heritable. BMI, Sex, and Age were frequently identified as being significant contributors to the variance of the measures. * The AD analysis resulted in warnings of high kurtosis, something known to affect heritability estimates. Despite transforming the variables using the inverse-log function, this could not be resolved. Therefore, the numbers given for the AD analyses should be interpreted with some caution. FA = Fractional Anisotropy, MD = Mean diffusivity, AD = Axial diffusivity, RD = Radial diffusivity.

| **Tract and Measure** | **Heritability** | **Significant Covariates** |
| --- | --- | --- |
| **Subic - BNST with Fornix (Mean FA)** | $H^2r = 0.51, p < 0.0001$ | Sex (females < FA, $p < 0.0001$), All covars together accounted for 7% of the variance. |
| **Subic - BNST excluding Fornix (Mean FA)** | $H^2r = 0.53, p < 0.0001$ | Sex (females < FA, $p < 0.0001$). All covars together accounted for 8% of the variance. |
| **Subic - BNST with Fornix (Mean MD)** | $H^2r = 0.66, p < 0.0001$ | BMI (< MD, $p < 0.0001$), Age (< MD, $p = 0.002$), Sex (females < MD) $p < 0.0001$). All covars together accounted for 30% of the variance. |
| **Subic - BNST excluding Fornix (Mean MD)** | $H^2r = 0.62, p < 0.0001$ | BMI (< MD, $p < 0.0001$), Age (< MD, $p = 0.005$), Sex (females < MD, $p < 0.0001$). All covars together accounted for 26% of the variance. |
| **Subic - BNST with Fornix (Mean AD)** | $H^2r = 0.86^*, p < 0.0001$ | BMI (< AD, $p < 0.0001$), Age (< AD, $p = 0.005$), Sex (females < AD $p < 0.0001$). All covars together accounted for 48% of the variance. * |
| **Subic - BNST excluding Fornix (Mean AD)** | $H^2r = 0.81^*, p < 0.0001$ | BMI (< AD, $p < 0.0001$), Age (< AD, $p = 0.006$), Sex (females < AD, $p < 0.0001$). All covars together accounted for 49% of the variance. * |
| **Subic - BNST with Fornix (Mean RD)** | $H^2r = 0.54, p < 0.0001$ | BMI (< RD, $p < 0.0001$), Age (< RD, $p = 0.001$), Sex (females < RD, $p < 0.0005$). All covars together accounted for 10% of the variance. |
| **Subic - BNST excluding Fornix (Mean RD)** | $H^2r = 0.51, p < 0.0001$ | BMI (< RD, $p < 0.0001$), Age (< RD, $p = 0.02$), Sex (females < RD, $p < 0.001$). All covars together accounted for 8% of the variance. |

## 4. Discussion

### 4.1 Summary of Results

We utilised autoradiographic histological tract-tracing analysis and dMRI computational tract-tracing (tractography) to delineate the structural connectivity between the subiculum complex and BNST in macaques and humans. Autoradiographic tracing revealed evidence for a direct monosynaptic projection from the hippocampal formation to the BNST, resulting in a discrete area of terminal label alongside labelled fibres. Follow-up analysis of a different macaque sample using probabilistic dMRI tractography revealed complementary in vivo evidence for a structural connection between the two regions. A similar pattern of results was found using probabilistic dMRI tractography in a large human sample from the HCP. Examination of the tract-tracing results show that this connection depends primarily on the fornix, with the diffusion tractography data suggesting an additional connection through a pathway involving the amygdala. In our human sample, we tested the heritability of white-matter microstructure indices (FA, MD, RD, AD), within our tracts and report that all measures were moderately heritable. Finally, in our HCP sample, we examined the putative functional attributes of the identified pathway, finding no evidence for a phenotypic, environmental, or genetic association between tract microstructure and components reflecting dispositional negativity and alcohol use. We did, however, find significant microstructure associations with our covariates BMI, age, and sex.

### 4.2 Autoradiographic Tract-Tracing of a Direct Hippocampal-BNST Projection in the Macaque

Results from our macaque autoradiographic tract-tracing study indicate that there is a direct, fornical projection from the subiculum to the BNST. In contrast, no BNST label was seen when the subiculum was not involved in the tracer injection, nor in those cases where the fornix was sectioned prior to tracer injection. In the case with the largest hippocampal injection (ACy28) this projection was especially evident, principally terminating in the medial division of the anterior part of the BNST. Other cases showed evidence of termination in the same area, although it was much lighter, potentially reflecting the relative sizes of the injections. This evidence for a direct projection via the fornix accords with previous NHP studies using degeneration methods in squirrel monkeys (Morrison & Poletti, 1980; Poletti & Creswell, 1977). Aside from case ACy28, the projections observed in the present study appeared modest, suggesting a potential difference from rodent brains (Ding, 2013; Ding et al., 2020). It is, however, the case that the rostral hippocampal injections (ACy12, ACy14) were largely confined within the subiculum and prosubiculum. In contrast, the large caudal hippocampal injection (ACy28) not only more completely filled the prosubiculum and subiculum, but also involved the CA fields and dentate gyrus. Consequently, the possibility remains of further hippocampal projections to the BNST from other areas within the hippocampal formation.

It is also possible that more extensive projections would have been seen if the tracer injections had included the presubiculum, as in the rat this area also projects to the BNST (Howell et al., 1991). A further reason for the modest connectivity seen in our NHP autoradiographic sample may be that subiculum influences on the BNST are mediated via the amygdala to a larger extent than in rodents. Numerous authors have emphasised the importance of this subiculum – amygdala - BNST route in rodents and in NHPs (Cullinan et al., 1993; Ding et al., 2020; Herman et al., 2020; Morrison & Poletti, 1980; Poletti et al., 1984; Poletti & Sujatanond, 1980), but if in primates this route is solely indirect, i.e., polysynaptic, then this connection would not be visible with the anterograde tracing method used in the present study.

Previous studies in both rats and squirrel monkeys have indicated that a significant proportion of hippocampal to BNST inputs are sent via non-fornical routes (Cullinan et al., 1993; Morrison & Poletti, 1980, 1980; Poletti et al., 1984). Indeed, following an electrophysiological study of hippocampal - amygdala - basal-forebrain connectivity in rats, the authors suggested that unit activity response latency times were indicative of the amygdala serving as a relay between the subiculum and the basal forebrain (Morrison & Poletti, 1980; Poletti et al., 1984). Therefore, given these previous findings, and that we only see fornical inputs with our monosynaptic tracer, it seems likely that in primates any substantial non-fornical influences will prove to be polysynaptic. It is notable that rodents have more monosynaptic non-fornical hippocampal efferent routes to other subcortical targets (Dillingham et al., 2015; Meibach & Siegel, 1975, 1977) than those described in non-human primates, so the BNST may be a further case of this.

### 4.3 Macaque Tractography Evidence for Subiculum Complex – BNST Structural Connectivity

Results from our macaque diffusion tractography analysis revealed evidence for connectivity between the subiculum complex and the BNST. Comparing the proportion of connectivity to other ROIs, results suggest that the BNST is second in terms of connection proportion to the ATN, and above that of the GPe and NAc. The difference with the ATN, which numerous studies have shown to have dense connectivity with the subiculum (Aggleton, 1986; Aggleton & Christiansen, 2015), was statistically significant. However, the BNST’s difference with either the negative comparison ROI (GPe), or the other positive comparison ROI (NAc) (see section 4.6) was not significant following Bonferroni correction. This likely reflects low sensitivity, given that our sample size was only five following outlier removal. Viewing the subiculum ROI – BNST tractography output image (Figure 6, also uploaded to Neurovault at https://identifiers.org/neurovault.image:682610) we see that the reconstructed tracts traversed dorsally via the fornix or ventrally via the amygdala, likely through the amygdalofugal pathway. This finding of a pathway involving the amygdala is consistent with previous research in squirrel monkeys and rodents (Cullinan et al., 1993; Herman et al., 2020; Morrison & Poletti, 1980, 1980; Poletti et al., 1984). The limitations of tractography analysis mean that our results cannot differentiate between mono- and poly-synaptic pathways (Campbell & Pike, 2014), therefore, that we see an amygdala pathway with tractography but not autoradiography is consistent with the hypothesis that this connection is polysynaptic in nature (Morrison & Poletti, 1980).

### 4.4 HCP Probabilistic Tractography Findings

Our HCP results suggest that the subiculum complex has substantial connectivity to the BNST in humans, with only the ATN receiving a higher proportion of streamlines of our tested ROIs. The reconstructed tracts appear to follow what would be expected given the rodent literature (Herman et al., 2020) and our macaque tractography results, with connectivity passing antero-dorsally through the fimbria/fornix and through another antero-ventral route via the amygdala, again supporting previous findings in the rat and squirrel monkey (Cullinan et al., 1993; Herman et al., 2020; Morrison & Poletti, 1980, 1980; Poletti et al., 1984). At least part of this amygdala route appeared consistent with the amygdalofugal pathway, as previously reported in rats (Poletti et al., 1984).

When testing whether excluding (via masking) the fornix had a significant effect on the number of streamlines between the subiculum ROI and the BNST, we found that this did indeed make a statistically significant difference. However, given that the fornix is the main output region of the hippocampus, the key insight garnered from this test was that more than half of the reconstructed streamlines remained. When viewing the fornix-excluded tractography output image (Figure 7, uploaded to Neurovault at https://identifiers.org/neurovault.image:682612), it is clear that some streamlines around the fornix structure remained. Part of this could be explained by imprecise registration of the NOT fornix mask to the subject space and individual differences in anatomy meaning that the NOT mask does not capture all the fornix pathway. Also likely, is that a substantial number of the remaining streamlines around the fornix NOT mask are from the stria terminalis pathway, which runs just laterally to the columns of the fornix and is the primary input to the BNST from the amygdala (Alheid, 2009; Alheid et al., 1998; Oler et al., 2017).

### 4.5 Macaque and Human Diffusion Tractography Results are Strikingly Similar

The similarity of the macaque tractography results to those in our human sample suggests that macaque monkeys can be a useful model for further investigation of this connection in humans. Macaques are much closer to humans in evolutionary terms compared to rodents (Murray et al., 2017) and monkey experiments have already demonstrated their utility in other areas of BNST/ stress research (Fox et al., 2018; Mars et al., 2018; Oler et al., 2017). Thus, contingent on larger studies verifying our findings, our results demonstrate that key pathways from the subiculum complex to the BNST can be detected using tractography methods and that the results are very similar between macaques and humans.

### 4.6 Human Subiculum Complex – BNST Tract Microstructure Heritability and Potential Functional Attributes

Microstructure metrics extracted from our subiculum ROI to BNST tracts were all shown to be moderately to largely heritable, ranging from a h^2^ of .41 to .69. This is in line with other twin-based heritability studies, which have demonstrated that white matter microstructure is influenced to a large degree by genetic factors (Gustavson et al., 2019; Lee et al., 2017; Vuoksimaa et al., 2017). We did not find evidence for any of the microstructure indices being significantly associated, genetically or otherwise, with our two components representing dispositional negativity traits and alcohol use.

Many rodent studies implicate the subiculum – BNST - PVN connection as being key to stress-related processing, with aberrant stress-processing having been proposed as a key mechanism linked to dispositional negativity (Herman et al., 2020; Hur et al., 2019; Radley & Johnson, 2018; Shackman et al., 2016). Structural and functional alterations of the hippocampus and/or BNST have previously been linked to stress-related traits and disorders (Avery et al., 2016; Hur et al., 2019). Although we could find no examples of studies investigating subiculum - BNST tract microstructure, a small number of studies have examined dMRI associations in the fornix with stress-related traits; with varying results (Barnea-Goraly et al., 2009; Benear et al., 2020; Modi et al., 2013; Yu et al., 2017).

One reason that we did not observe an association may be because our sample consists of a screened, relatively healthy population. Evidence for BNST involvement in dispositional negativity or stress-related traits generally comes from studies of clinical populations, or during experiments when states such as fear/anxiety have been induced (Brinkmann et al., 2018; Klumpers et al., 2017; Mobbs et al., 2010; Pedersen et al., 2020). In addition, some researchers have proposed that human self-report emotions should be considered as distinct from the sub-cortical and physiological processes often studied in animal models (LeDoux & Hofmann, 2018). If correct, this would render association between our self-report questionnaire data and our sub-cortical structural connection unlikely. Alternatively, a large-scale study (n=559) recently reported that stress-reactivity may be less important for traits related to dispositional negativity than previously thought (Mineka et al., 2020).

Both the hippocampus and the BNST have been associated with alcohol use, with the BNST being specifically implicated in alcohol-related alterations to stress-processing (Bach et al., 2021; Volkow et al., 2016). One recent small-scale study reported a greater number of streamlines between the hippocampus and the BNST in early abstinence women than in control women, a finding which was not seen in men (Flook et al., 2021). Although interesting, assessing streamlines in this way should generally be avoided in favour of microstructure measures, as individual differences in streamline variation are difficult to interpret (Jones et al., 2013). Whilst not specific to the BNST, white matter abnormalities of the fornix in adults with heavy drinking and alcohol use disorder have been reported (Cardenas et al., 2013), although the diffusion MRI literature in general with regards to correlates of substance abuse is somewhat inconsistent (Benear et al., 2020). Similar to our dispositional negativity component, a reason we did not find such an association could be that our sample consisted of a screened, relatively healthy population (median drinks consumed per week being two), which likely reduced our chances of finding any association (Berry et al., 2020). Studies that have found associations between alcohol use and white matter metrics have generally been conducted either with clinical populations, or following experimental manipulations (Campbell et al., 2019; Cardenas et al., 2013; Flook et al., 2021).

We report an effect of our covariates BMI, age, and sex on nearly all the microstructure measures, on tracts both including and excluding the fornix (Table 4). These associations were expected given that all of these factors have been previously associated with global alterations in white-matter microstructure (Dekkers et al., 2019; Lawrence et al., 2021; van Hemmen et al., 2017; Xiong & Mok, 2011; Yang et al., 2016). Due to these global effects, understanding microstructural relationships with these traits along specific tracts is a non-trivial problem that requires careful analysis. Specific investigation with regards to the associations of these factors with microstructural properties along the subiculum- BNST-PVN pathway would be informative, particularly as the BNST has been associated with feeding behaviours and is sexually dimorphic (Allen & Gorski, 1990; Lebow & Chen, 2016).

### 4.7 Limitations

An unexpected result was that the additional positive comparison region, the NAc, appeared to demonstrate little evidence of subiculum connectivity via probabilistic tractography in both macaques and humans. This connection was detected in our tract-tracing analysis and has been widely described elsewhere (Aggleton & Christiansen, 2015). This result is likely due to the limitations of tractography, which can struggle to resolve crossing and fanning fibre populations, even at relatively high image resolution (Schilling et al., 2017). The NAc is connected to the subiculum via the fornix. The fornix bifurcates at the anterior commissure into the postcommissural and precommissural fornix, with part of the precommissural fornix travelling anteriorly and ventrally to terminate in the nucleus accumbens, with adjacent fibres reaching frontal areas (Aggleton & Christiansen, 2015; Brog et al., 1993; Friedman et al., 2002; Groenewegen et al., 1987). It may be that due to the nature of this bifurcating tract, the extent of the precommissural connection is underestimated by probabilistic tractography. This specific problem has been noted elsewhere (Brown et al., 2017). Another limitation is that the tensor-derived white matter microstructure indices used in our sample are only indirect measures inferred from water diffusion, and thus do not reflect more specific properties of white matter microstructure (Afzali et al., 2021). More biologically meaningful measures, such as those derived from biophysical models e.g., CHARMED (Assaf & Basser, 2005), or NODDI (Zhang et al., 2012), may demonstrate evidence for specific microstructural features that are related to our phenotypes of interest, although DTI is more sensitive to individual differences than more specific microstructure measures (De Santis et al., 2014). A further limitation involves our use of self-report questionnaire data, which may only be indirectly (if at all) related to subcortical stress-processing mechanisms (LeDoux & Hofmann, 2018; Mineka et al., 2020).

### 4.8 Conclusions and Future Directions

We used a multi-method, cross-species approach (Folloni et al., 2019) to demonstrate that a key connection identified in rodent stress research, between the subiculum complex and BNST, is present in macaques and humans. We show that this connection has a substantial fornical element, with our diffusion tractography results suggesting an additional route via the amygdala, which may be more polysynaptic in nature than the monosynaptic pathways described in rodent research (Cullinan et al., 1993; Poletti et al., 1984). As such, further research is needed to indicate to what extent the amygdala plays a role as an intermediary between the subiculum and BNST in primates. Further refinements should include using tractography to compare the BNST connectivity of subiculum sub-regions (e.g., the prosubiculum, subiculum, presubiculum, parasubiculum). Recent rodent research has implicated the ventral (anterior in primates) subiculum, or prosubiculum, in connections to the BNST, with suggestions that connectivity differences are distributed along a ventral-dorsal gradient (Ding, 2013; Ding et al., 2020). Given that most of this research has taken place in rats, and given the potential species differences reported here, the nature of subicular subregion projections to the BNST should be further investigated in primates. Methods to achieve this include the application of advanced ultra-high field MRI techniques specifically developed to analyse hippocampal subfields (e.g., Hodgetts et al., 2017), or ultra- high resolution dMRI applied to post-mortem macaque brains (e.g., Sébille et al., 2019). Although we did not find any associations with our principal components relating to dispositional negativity or alcohol use, differences may yet be found when studying clinical populations (e.g., Cardenas et al., 2013), the impact of chronic early life stress (Petrican et al., 2021), or physiological and behavioural/cognitive biomarkers of stress-responding (e.g., Allen et al., 2017). In general, though we describe in humans and macaques the existence of a subiculum-BNST connection previously identified in rats, the appreciation of potential species differences demonstrated here has implications for the generalisability of rodent research to the understanding of human stress-related functional anatomy and its disorders.

## Acknowledgements

This project was funded by a Wellcome Trust PhD studentship awarded to Samuel Berry, grant reference: 215194/Z/19/Z. Human data were provided [in part] by the Human Connectome Project, WU-Minn Consortium (Principal Investigators: David Van Essen and Kamil Ugurbil; 1U54MH091657) funded by the 16 NIH Institutes and Centers that support the NIH Blueprint for Neuroscience Research; and by the McDonnell Center for Systems Neuroscience at Washington University. Our thanks to Shen K, Bezgin G, Schirner M, Ritter P, Everling S, McIntosh AR & OpenNEURO ([Dataset] doi: 10.18112/openneuro.ds001875.v1.0.3) for the collection and hosting of the non-human primate neuroimaging data. Special thanks to Christopher Dillingham at Cardiff University for producing the bright field, dark field, and overlay images (Figure 4).

