## supplementary material for "Subiculum – BNST Structural Connectivity in Humans and Macaques"

### HCP Demographics

| **Family Size** | **Count** |
| --- | --- |
| 1 | 74 |
| 2 | 162 |
| 3 | 178 |
| 4 | 35 |
| 5 | 3 |
| 6 | 1 |

| **Ethnicity** | **Count** |
| --- | --- |
| White | 734 |
| Black or African American | 176 |
| Hispanic/ Latino | 95 |
| Asian/ Nat. Hawaiian/ Other Pacific Is. | 61 |
| More than one | 22 |
| Unknown or Not Reported | 3 |
| Am. Indian/ Alaskan Nat. | 2 |

| **Age Group** | **Count** |
| --- | --- |
| 22-25 | 227 |
| 26-30 | 477 |
| 31-35 | 377 |
| 36+ | 12 |

### Probtrackx2 options

Macaques

probtrackx2 –network -x -l --onewaycondition -c 0.2 -S 2000 --steplength=0.2 -P 5000 --fibthresh=0.01 --distthresh=0.0 --sampvox=0.0 –forcedir --pd --opd -s merged -m nodif_brain_mask.nii.gz

Humans

probtrackx2 --network -x -l --onewaycondition -c 0.2 -S 2000 --steplength=0.5 -P 5000 --fibthresh=0.01 --distthresh=0.0 --sampvox=0.0 --xfm= --invxfm=--forcedir --opd -s -m nodif_brain_mask

### Bivariate Solarius Outputs

| **Tract and Measure** | **PC1 - RhoP** | **PC1 RhoE** | **PC1 - RhoG** | **PC2 - RhoP** | **PC2 - RhoE** | **PC2 - RhoG** |
| --- | --- | --- | --- | --- | --- | --- |
| **Subic - BNST with Fornix (Mean FA)** | RhoP = 0.02 , p = 0.6 | RhoE = 0.05(SE 0.08), p = 0.51 | RhoG = -0.06 (SE 0.19), p = 0.75 | RhoP = -0.05, p = 0.10 | RhoE = -0.08 (SE 0.08), p = 0.32 | RhoG = -0.003 (SE 0.18), p = 0.98 |
| **Subic - BNST excluding Fornix (Mean FA)** | RhoP = 0.004, p = 0.89 | RhoE = 0.05 (SE 0.09), p = 0.58 | RhoG = -0.08 (SE 0.19), p = 0.66 | RhoP = - 0.03, p = 0.39 | RhoE = 0.02 (SE 0.08), p = 0.81 | RhoG = -0.13 (SE 0.18), p = 0.47 |
| **Subic - BNST with Fornix (Mean MD)** | RhoP = -0.05, p = 0.12 | RhoE = 0.004 (SE 0.9), p = 0.96 | RhoG = -0.16 (SE 0.18), p = 0.34 | RhoP = 0.01, p = 0.83 | RhoE = 0.004 (SE 0.09), p = 0.96 | RhoG = 0.01 (SE 0.17), p = 0.94 |
| **Subic - BNST excluding Fornix (Mean MD)** | RhoP = -0.03, p = 0.31 | RhoE = -0.02 (SE 0.09), p = 0.81 | RhoG = -0.06 (SE 0.19), p = 0.74 | RhoP =-0.002 p = 0.94 | RhoE = - 0.05 (SE 0.08), p = 0.52 | RhoG = 0.10 (SE 0.18), p = 0.58 |
| **Subic - BNST with Fornix (Mean AD)** | RhoP = -0.05. p = 0.11 | RhoE = 0.06 (SE 0.11), p = 0.55 | RhoG = - 0.23 (SE 0.17), p = 0.17 | RhoP = - 0.02, p = 0.52 | RhoE = -0.06 (SE 0.10), p = 0.56 | RhoG = 0.02 (SE 0.16), p = 0.89 |
| **Subic - BNST excluding Fornix (Mean AD)** | RhoP = -0.05, p = 0.09 | RhoE = 0.01 (SE 0.10), p = 0.93 | RhoG = -0.17 (SE 0.17), p = 0.32 | RhoP = - 0.01, p = 0.85 | RhoE = -0.02 (SE 0.09), p = 0.86 | RhoG = 0.01 (SE 0.16), p = 0.96 |
| **Subic - BNST with Fornix (Mean RD)** | RhoP = -0.03, p = 0.33 | RhoE = -0.01 (SE 0.08), p = 0.86 | RhoG = -0.07 (SE 0.20), p = 0.71 | RhoP = 0.01, p = 0.65 | RhoE = 0.02 (SE 0.8), p = 0.82 | RhoG = 0.007 (SE 0.19), p = 0.97 |
| **Subic - BNST excluding Fornix (Mean RD)** | RhoP = -0.01, p = 0.66 | RhoE = -0.03 (SE 0.08), p = 0.70 | RhoG = 0.03 (SE 0.21), p = 0.90 | RhoP = - 0.01, p = 0.82 | RhoE = -0.06 (SE 0.08), p = 0.47 | RhoG = 0.11 (SE 0.19), p = 0.58 |

Supplementary Table 1

Bivariate heritability analyses between each extracted DTI measure and each principal component revealed no phenotypic, environmental, or genetic associations. PC1 represented traits related to dispositional negativity, whereas PC2 reflected measures of alcohol use.
